# Ternary Neurexin-T178-PTPR complexes represent a presynaptic core-module of neuronal synapse organization

**DOI:** 10.1101/2024.07.16.603715

**Authors:** Spyros Thivaios, Jochen Schwenk, Aline Brechet, Sami Boudkkazi, Nithya Sethumadhavan, Phil Henneken, Eriko Miura, Ayumi Hayashi, Maciej K. Kocylowski, Alexander Haupt, Debora Kaminski, Dietmar Schreiner, Jean-Baptiste van den Broucke, Akos Kulik, Uwe Schulte, Fredrik H. Sterky, Michisuke Yuzaki, Peter Scheiffele, Bernd Fakler

## Abstract

The organization of cell-cell contacts is fundamental for multi-cellular life and operation of organs. Synapses, prototypic contact sites for neuronal communication, are key to brain function and work over the last decades identified multiple synaptic cell adhesion molecules (sCAMs) that drive their organization. Whether these sCAMs operate independently or in coordination through yet unknown linker proteins remained elusive. Here, we used a systematic large-scale multi-epitope affinity-purification approach combined with quantitative mass spectrometry and immuno-EM to comprehensively map trans-synaptic protein networks in the mouse brain. We discover a presynaptic core-module assembled from the two major sCAM families, Neurexins1-3 and LAR-type receptor protein tyrosine phosphatases (PTPRD,S,F), and the previously uncharacterized tetraspanin proteins T178A, B. These ternary Neurexin-T178-PTPR complexes form through their trans-membrane domains and assemble during biogenesis in the ER. Loss of T178B results in module dissociation, strong reduction of LAR-PTPRs and re-distribution of synaptic Neurexins. At synapses, the Neurexin-T178-PTPR module recruits stable and extended trans-synaptic protein networks with defined pre- and post-synaptic partners and secreted extracellular linkers. The network architecture robustly interlinks the distinct functional modules/machineries of the presynaptic active zone and establishes tight associations with XKR-type lipid scramblases and postsynaptic GABAergic and glutamatergic neurotransmitter receptors. Our data identify a universal presynaptic core-module for synaptic adhesion and trans-synaptic signaling in the mammalian brain.

## Introduction

Neuronal synapses are highly specialized units for cell-cell communication that underlie the transmission and storage of information fundamental for proper operation of the brain and its circuits. The functional properties of synapses display profound variability across the CNS and are to a significant extent instructed by synaptic cell adhesion molecules (sCAM). These sCAMs, mostly single pass (or type I) transmembrane proteins, drive formation and structural organization of neuronal synapses and regulate their stability and context-dependent dynamics (Aoto et al., 2015; de Wit and Ghosh, 2016; Gomez et al., 2021; Krueger-Burg et al., 2017; Uemura et al., 2010).

While individual sCAMs were found sufficient to induce neuronal synapse formation in culture systems (Biederer et al., 2017; Graf et al., 2004; Scheiffele et al., 2000), their deletion by targeted gene-knockout only moderately reduced synaptogenesis in the mammalian brain (Anderson et al., 2015; Chen et al., 2017; Emperador-Melero et al., 2021; Horn et al., 2012; Missler et al., 2003; Takahashi and Craig, 2013; Varoqueaux et al., 2006). This resilience to genetic perturbation appeared to argue for a high degree of redundancy, and led to the postulation of two distinct models (for explanation): (1) Multiple adhesion systems act independently or, interlinked by yet unknown adapters, in concerted fashion, or, alternatively, (2) there is a main essential synapse-organizing system that remains to be identified.

Among the currently known sCAMs, two families of proteins stand out based on their ubiquitous and highly abundant expression at central synapses: Neurexins (NRX) and the leukocyte common antigen-related receptor protein tyrosine phosphatases (LAR-type PTPRs). Both of these sCAM families of presynaptic type I transmembrane proteins comprise three distinct members, NRX1-3 and PTPRD, S and F, that display complex molecular appearances as a result of (marked) genetic variations/alternative splicing and interaction(s) with varieties of partner proteins. Thus, NRX1-3 use alternative promotors to generate long (α-neurexin) and short (β-neurexin) isoforms (Hauser et al., 2022; Schreiner et al., 2014; Sudhof, 2017; Ushkaryov et al., 1992); in addition, for *Nrxn1*, a very short γ-neurexin isoform is expressed from a third alternative promoter (Sterky et al., 2017; Yan et al., 2015) (**Figure 1A**). NRXs contribute to presynaptic active zone assembly, impact Ca^2+^-dependent neurotransmitter release, and instruct trans-synaptic nano-alignments in mice, flies and worms via interaction with multiple partners including neuroligins (NLGNs), leucine-rich repeat transmembrane neuronal proteins (LRRTMs) and cerebellins (CBLNs). LAR-PTPRs exhibit evolutionarily conserved cytoplasmic interactions with key regulators of active zone assembly, most prominently complexes of SYD-1 and SYD-2/Liprin-α (LIPA) proteins (Dai et al., 2006; Marco de la Cruz et al., 2024; Muhammad et al., 2015; Owald et al., 2012; Wentzel et al., 2013; Wong et al., 2018), thus tethering transmitter vesicles to the active zone (termed priming complexes (Emperador-Melero et al., 2024)). Notably, recent work reported severely impaired synapse development in cerebellar circuits upon simultaneous deletion of three NRX and three LAR-PTPR proteins and led to the suggestion that their ‘combinatorial’ expression may ensure common, though mutually independent action(s)(Sclip and Sudhof, 2023). However, understanding the organizing principles and relation of the multiple sCAMs expressed at individual synapses remains a major unresolved question.

**Figure 1.**
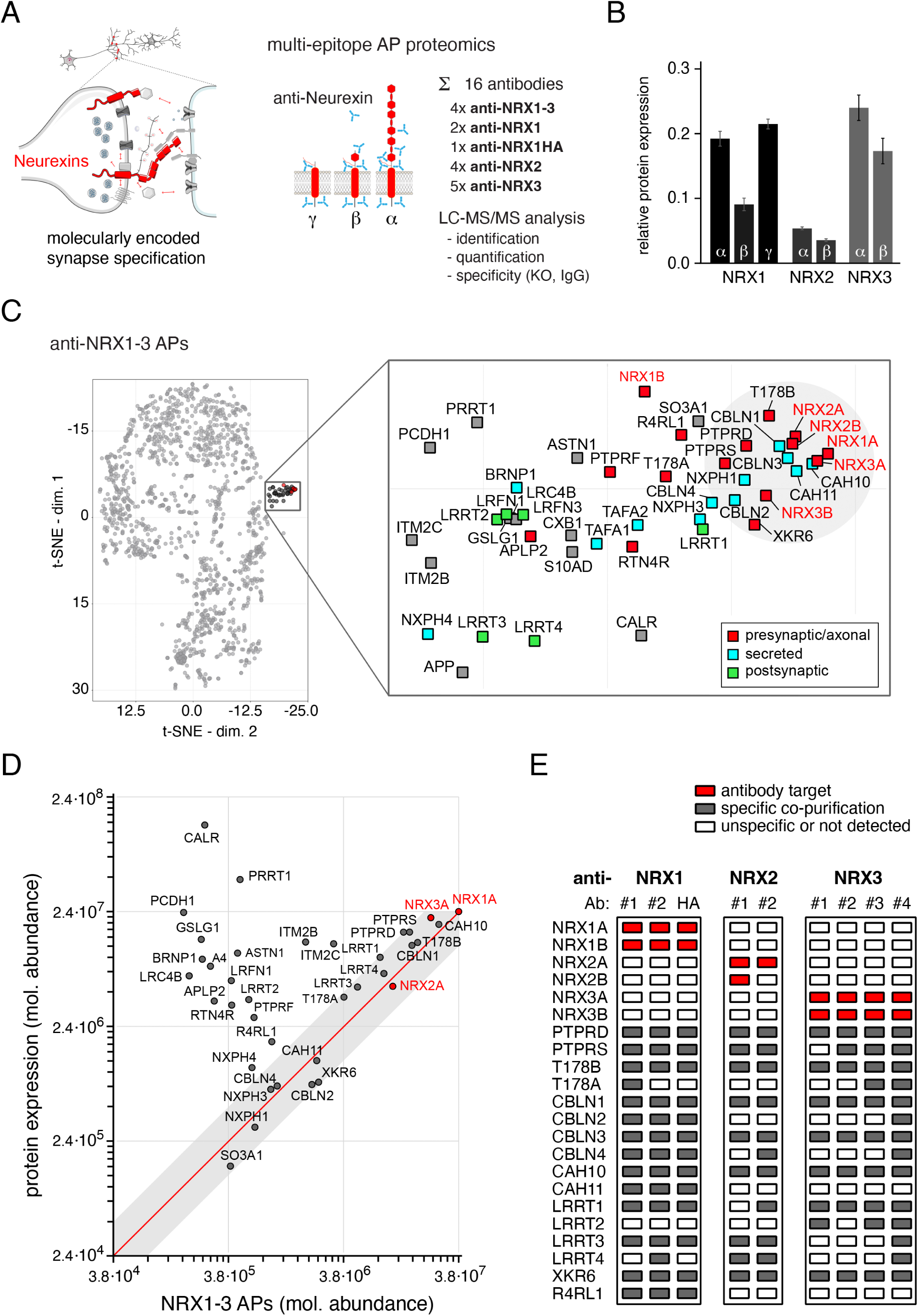
meAP-MS analysis of neurexin-associated protein assemblies. (A), Proteomic meAP-MS approach for quantitative analysis of NRX-associated trans-synaptic protein complexes. (B), MS-derived protein abundance of all isoforms of the NRX paralogues in plasma membrane-enriched fractions from adult mouse brain (technical details in **Supplementary Figure 1**). Data are mean ± SD of 4 brain samples. (C), t-SNE plot based on target normalized ratio (tnR)-values of all proteins (grey dots; Methods) identified in eight APs with the four distinct anti-NRX1-3 ABs with preimmunization IgGs and antiserum as negative controls and NRX1A as a reference for normalization. Right panel, extension of the framed area around the target proteins (NRX1-3) highlighting a cluster of the consistently co-purified proteins of mainly pre-, post- and trans-synaptic origin as indicated by the color-code. (D), MS-derived molecular abundance values determined for the constituents of the NRX-cluster in (C) in (untreated) mouse brain membrane fractions plotted against the molecular abundance values determined in anti-NRX1-3 AP eluates. Proteins with complete binding to the NRX-target(s) are located on the diagonal in red. Area in grey defines the confidence-interval of MS-quantification. Data are mean values from four mouse brains and eluates of four independent APs with two different ABs. (E), Table summarizing a selected set of proteins specifically co-purified in APs with the indicated isoform-specific anti-NRX antibodies. Specificity was determined with target-KO (NRX1 for HA-tagged NRX1, NRX3) and IgG controls. Note that PTPRD,S and T178B are interactors of all NRX isoforms.

Here, we used an unbiased quantitative analysis of native protein assemblies in synaptic membranes by multi-epitope affinity-purifications (meAPs) combined with high-resolution mass spectrometry (nano-LC MS/MS) to uncover ternary complexes of NRXs, LAR-PTPRs and the tetraspanins T178A,B as a broadly expressed core-module for organization of signaling entities in the pre-synapse and for establishment of high-affinity associations with postsynaptic receptor and adhesion systems.

Added Note:

Proteins will be referred to by the acronyms used in the UniProt/SwissProt database throughout, descriptive names or gene names may be taken from Table 1 or text.

**Table 1.**
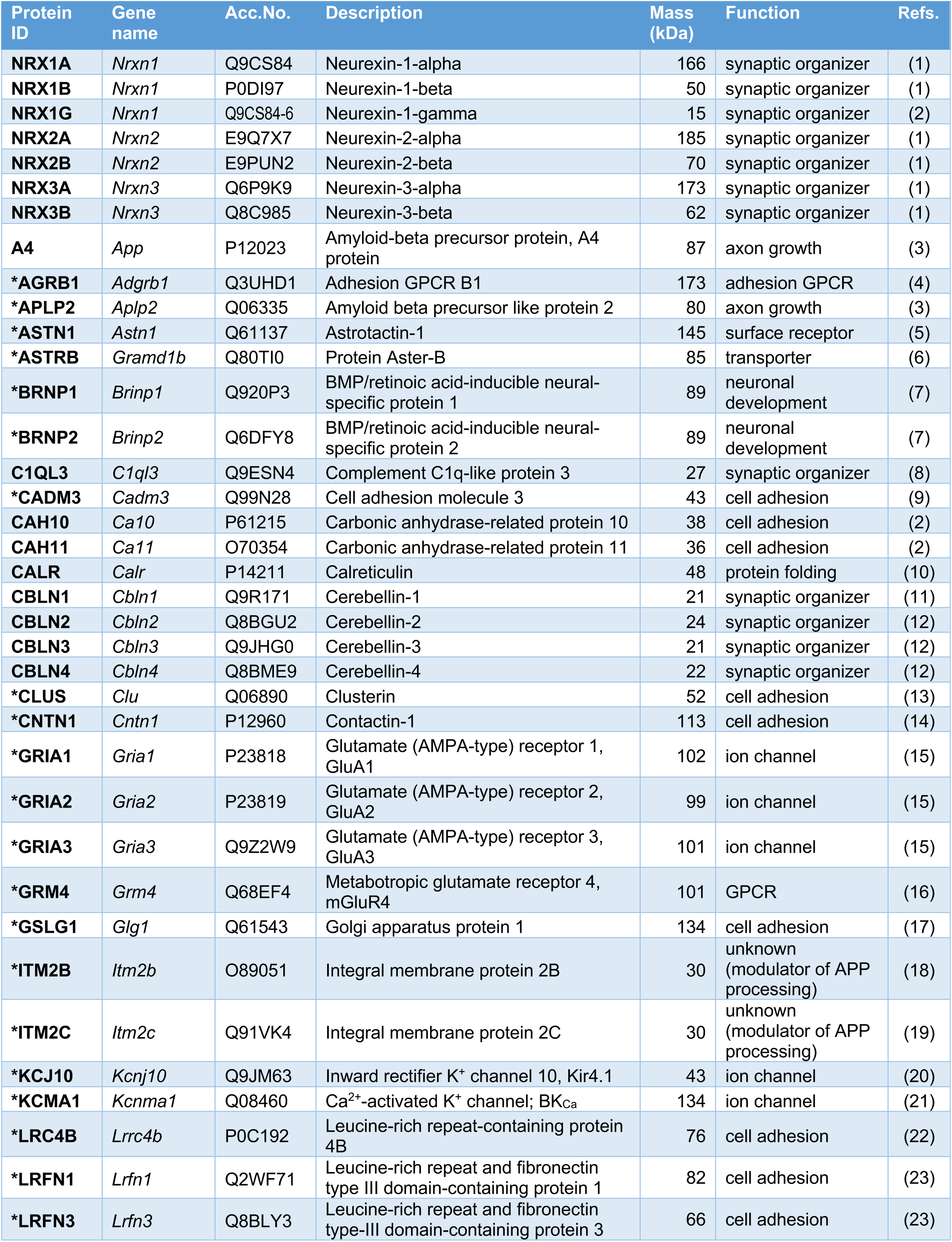
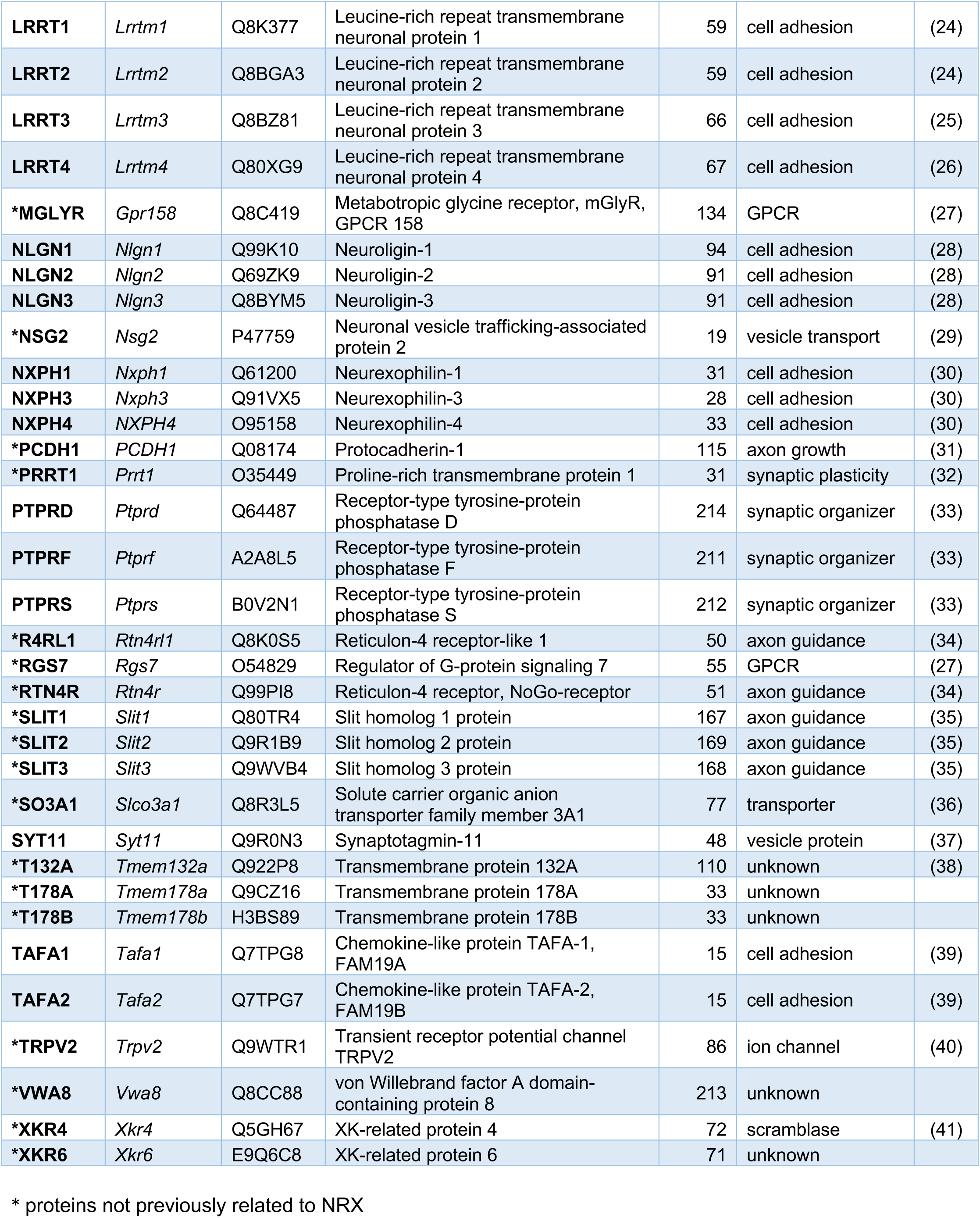
Interactome of native NRX1-3. (identified by meAP-MS with anti-NRX1-3 ABs) All proteins identified by quantitative MS-analysis in the APs with the four different anti-NRX1-3 ABs were evaluated by determination of tnR-values (Methods, text) using target-unrelated ABs and pre-immunization IgGs as negative controls. All proteins with tnR-values >0.25 in APs with a least two of the four ABs were considered as specific interactors of NRXs; the entirety of these specific interactors is referred to as NRX-interactome. Accordingly, the NRX-interactome comprises additional constituents compared to the t-SNE cluster (**Figure 1c**) of proteins consistently retrieved in APs with all four NRX1-3 ABs.

## Results

### Synaptic adhesion molecules are embedded into extended protein networks

Previous studies probed interactions and functions of individual sCAMs or sCAM families. To obtain a detailed map of sCAM interactomes in native tissue, we developed a comprehensive high-resolution proteomics approach combining multi-epitope affinity-purifications with quantitative mass spectrometry (meAP-MS; Methods) for an array of synaptic proteins. Initially, we leveraged a set of 16 AP-validated antibodies (ABs, **Supplementary Table 1**) targeting the primary NRXs via epitopes that are either isoform-specific (anti-NRX1/anti-NRX1HA, anti-NRX2, anti-NRX3) or present in all NRXs (anti-NRX1-3 or panNRX) and that are located at various positions along the primary sequences (**Figure 1A**, left, **Supplementary Table 1**, Methods). The input for the meAPs were membrane-fractions from adult mouse brains (see Methods) which were treated with the detergent CL-91 established in previous work for effective solubilization of protein assemblies in synapses (Boudkkazi et al., 2023; Muller et al., 2010; Schwenk et al., 2012; Schwenk et al., 2016). In fact, quantitative MS detected NRX1-3 in these plasma membrane-enriched fractions at distinct amounts with NRX1 and NRX3 being the most abundant (**Figure 1B**). Surprisingly, we found similar amounts for both the canonical long NRX1α and the only recently recognized short γ-isoform of NRX1, highlighting NRX1γ as a major NRX1 isoform in the mouse brain (**Figure 1B**, **Supplementary Figure 1**).

To uncover native NRX protein complexes, the four independent anti-NRX1-3 ABs were used in a first round of APs as target-specific ABs, together with pools of pre-immunization IgGs and pre-immunization sera as negative controls. All resulting AP-elutions were analyzed by high-resolution nano-LC MS/MS and evaluated by label-free quantification based on peak-volumes (PVs) and offering a linear dynamic range of about four orders of magnitude (Bildl et al., 2012; Boudkkazi et al., 2023; Schulte et al., 2023). The respective results showed effective purification of the NRX paralogues as indicated by large PVs and broad coverage of the primary sequences by MS/MS-identified peptides (**Supplementary Figure 2**). In addition to the NRX targets, the APs retrieved a number of additional proteins that were evaluated for their target-specific co-purification, as well as for consistency and direct comparability across the individual APs. The latter was done by determining for any protein target-normalized ratios (tnRs, (Kocylowski et al., 2022b), **Supplementary Figure 3A**) which is the relative abundance of a protein related to both, AP-background (minimal value derived from negative controls) and AP-target (maximal value; Methods, **Supplementary Figure 3A**). When analyzed by t-SNE (t-distributed stochastic neighbor embedding), the resulting plot of the tnR-values obtained in APs with the four anti-NRX1-3 ABs (**Supplementary Figure 3B**, **Table 1**) immediately identified a well-separated cluster of 45 proteins that tightly co-localized around the targets, thus indicating robust and consistent co-purification with the anti-NRX1-3 ABs in all APs (**Figure 1C**, **Supplementary Table 2**). Closer inspection uncovered notable features of these target-clustered proteins (**Figure 1C**, inset, **Table 1**): (1) They are components of the pre- and post-synaptic compartments, and the synaptic cleft, (2) most identified proteins are either single-pass trans-membrane or soluble/secreted proteins, and (3) several of them were established interaction partners of the NRX proteins, such as neurexophilins (NXPH1-4, (Missler and Sudhof, 1998)), cerebellins (CBLN1-3, (Matsuda and Yuzaki, 2011)), carbonic anhydrase-related (CAH) proteins 10, 11 (Sterky et al., 2017), chemokine-like proteins TAFA1,2 (Khalaj et al., 2020), amyloid-β precursor protein (A4 or APP, (Cvetkovska et al., 2022)), receptor protein tyrosine phosphatase S (PTPRS, (Roppongi et al., 2020)) or leucine-rich repeat transmembrane neuronal (LRRT) proteins 1-4 (de Wit et al., 2009; Roppongi et al., 2020; Siddiqui et al., 2010). In addition, the cluster contained a substantial number of proteins that were previously unknown to be associated with NRXs. These proteins include the tetraspanins T178A and T178B, the putative lipid scramblase XK-related protein 6 (XKR6), reticulon-4 receptor (RTN4R, also termed Nogo receptor) and reticulon-4 receptor like 1 (R4RL1), as well as amyloid-β precursor-like protein (APLP2), ITM2B, C (Schwenk et al., 2016), and the LAR-type receptor protein tyrosine phosphatases PTPRD and F. Interestingly, quantitative analysis of protein abundances in both source material (‘protein expression’) and elutions of NRX1-3APs showed that PTPRs D, S and the tetraspanin T178B are for most parts associated with the NRXs and that these co-assemblies contain the fast majority of expressed NRX proteins in the brain (**Figure 1D**, diagonal in red). Thus, NRXs, LAR-PTPRs and T178B are not only co-expressed across the entire brain (**Supplementary Figure 4**) but apparently occur in stable stoichiometric complexes, rather than as independent adhesive units. In addition, these analyses showed that the major portion of NRXs is covalently linked to CAH10, and interacts with the secreted CBLN1 protein (**Figure 1D**).

In a second round of APs, we next used the isoform-specific anti-NRX ABs and target-knockouts (**Supplementary Figure 5**) as stringent negative controls to uncover whether these novel interactors associate selectively with any of the NRX paralogues or whether they co-assemble with all primary NRX proteins. Importantly, we discovered that NRX1, 2, and 3 do not mutually co-purify (**Figure 1E**) indicating that they form separate molecular entities. While some of the aforementioned interactors, like CAH11, R4RL1 and LRRTM2, exhibited preferences for individual NRXs, others co-assembled with all three NRX paralogues (**Figure 1E**, **Supplementary Table 2**). Most prominently, PTPRs and T178B were effectively retrieved in all NRX APs, as was XKR6, albeit at markedly lower amounts (**Supplementary Figure 3B**).

Together, these meAP-MS data demonstrated that in the adult brain the three NRX paralogues form separate trans-synaptic modules with numerous interactors. However, all three NRXs co-assemble with T178B and LAR-PTPRs into previously unrecognized stoichiometric complexes.

### Neurexins form ternary core-modules with T178 and LAR-type PTPRs

The co-assembly of NRXs with T178 and LAR PTPRs was further investigated by native gel electrophoresis (BN-PAGE) and reverse APs. BN-PAGE analysis of brain membrane fractions indicated co-migration of NRX1-3A, LAR-PTPRs and T178B at an apparent molecular mass of roughly 400 kDa (**Figure 2A**, arrowhead) consistent with complex formation at an equimolar (or 1:1:1) stoichiometry based on predicted protein mass (**Table 1**). In addition, the BN-PAGE demonstrated assembly of the ternary complex with multiple additional partners/interactors as reflected by the extension of all three Western blot signals into the higher apparent mass range (**Figure 2A**). Orthogonal (‘reverse’) APs with ABs against both, PTPRS and T178B, robustly retrieved the NRX proteins together with multiple of their interaction partners as shown by the respective tnR-values (**Figure 2B**). Accordingly, the ternary NRX-T178-LAR-PTPRs complex may be considered a ‘core-module’ of larger assemblies whose constituents were consistently and robustly co-purified in the meAP experiments (**Figure 1C**, **E**, **Figure 2B**, **Table 1**).

**Figure 2.**
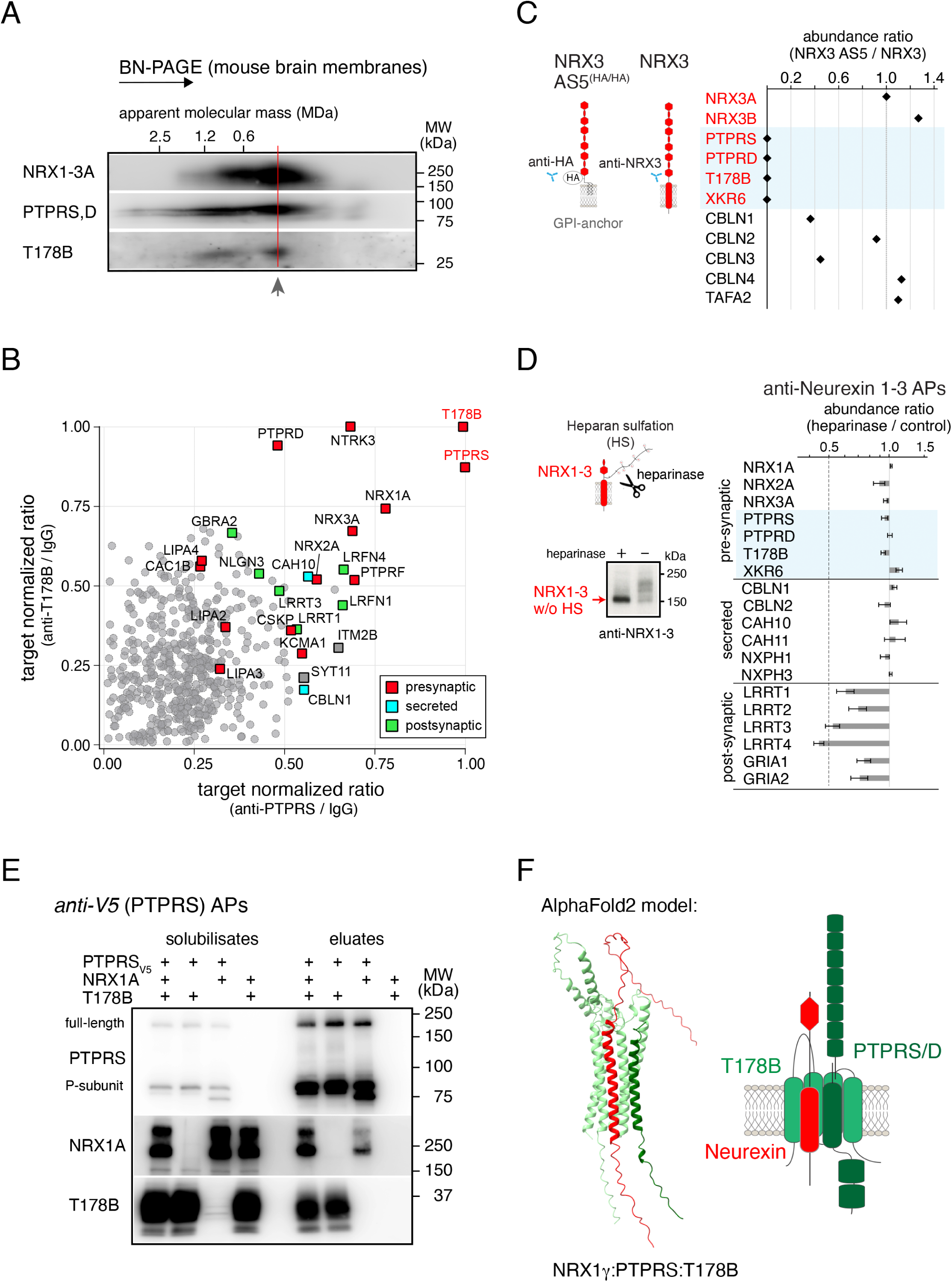
Ternary complex formation of Neurexins with LAR-type PTPRs and the tetraspanin T178B. (A), Native gel separation (BN-PAGE) of protein complexes in CL47-solubilized membrane fractions from adult mouse brain Western-probed for the indicated proteins. Note close co-migration of NRX1-3A, PTPRD/S and T178B as tripartite complex with an apparent molecular weight of approximately 400 kDa (arrowhead). Fainter signals extending to an apparent molecular mass of 1.2 MDa reflect their integration into larger complexes/entities. (B), tnR-plot comparing APs with ABs targeting T178B and PTPRS indicates robust mutual co-purification and enrichment of NRX1-3 and a variety of pre- and post-synaptic interactors. tnR-values were determined with IgGs as a reference. Color-code of subcellular localization is indicated. (C), Abundance ratio determined for proteins affinity-purified in anti-HA APs from HA-Neurexin3 AS5 (GPI-anchored splice variant of NRX3, inset) mice and in anti-NRX3 APs from wild-type mice. Note that GPI-anchored NRX3 complexes lack association with presynaptic PTPRD,S, T178B and XKR6, whereas binding of extracellular ligands was maintained. Experiments were done with two distinct HA and two NRX3 ABs, values are mean of two independent experiments. (D), Abundance ratio for a subset of proteins affinity-purified with anti-NRX1-3 from CL91-solubilized brain membrane fractions with and without enzymatic treatment with a mixture of heparinases (I, II, III); inset: Western-probed gel-separation indicating heparinase-mediated reduction in size of NRX3. Enzymatic activity strongly reduced co-purification of postsynaptic proteins (LRRTs, AMPAR subunits GRIA1, 2), while binding to presynaptic and extracellular/secreted interactors remained unchanged. Data are mean ± SEM of 3 experiments. (E), Reconstitution of ternary complexes assembled from PTPRS, NRX1A and T178B in cultured tsA cells. Specific co-purification of T178B and NRX1A with V5-tagged PTPRS in anti-V5 APs. The presence of T178B appears critical for stable complex formation with NRX1A and impacts maturation of the P-subunit of PTPRS. (F), Left panel, Structure of the membrane-adjacent region of the core-module predicted by AlphaFold2-Multimer with contact sites in the transmembrane plane. Input used for AlphaFold was NRX1γ (mouse, aa 1-139, full length), PTPRS (mouse, aa 1251-1360), T178B (human, 1-294, full length). Right panel, Scheme of the ternary complex with multiple sites/domains for protein-protein interactions on both the extracellular and the cytoplasmic side of the membrane.

Next, we investigated the structural determinant(s) for the assembly of the ternary core-module in native tissue by comparative APs (**Figure 2C**, **D**, insets) dissecting the role of the extracellular domain versus the transmembrane and intracellular stretches, and the significance of the heparan sulfate moieties (Zhang et al., 2018). We targeted the endogenous GPI-anchored NRX3 AS5 splice variant tagged through the knock-in of an HA-epitope (NRX3 AS5^(HA/HA)^, (Hauser et al., 2022)) and compared the recovery of the core-module components to APs for total NRX3 (**Figure 2C**). MS-based quantification indicated that assembly into the core-module is abolished for the splice-variant lacking the transmembrane domain, while binding of the soluble interactors (CBLNs or TAFAs) to the extracellular domains of NRX3 was largely preserved in the NRX3 AS5 proteoform (**Figure 2C**). In contrast, removal of heparan sulfation markedly decreased co-purification of post-synaptic interactors (including LRRTs and AMPA-type glutamate receptors (AMPARs)), but neither affected assembly of the core-module nor its association with soluble partner proteins (**Figure 2D**). Moreover, assembly of the core-module did not require any additional components as indicated by the robust complex formation observed upon heterologous expression of all three constituents (**Figure 2E**, **Supplementary Figure 6**). Interestingly, effective co-purification of PTPRS and NRX1A or NRX3A strongly depended on the presence of T178 in these AP-experiments, while weak interaction between NRX and PTPR was observed upon their pairwise expression in the absence of the tetraspanin (**Figure 2E**, **Supplementary Figure 6**). Likewise, binding of GPI-anchored NRX3A to PTPRS was weak and independent of T178B (**Supplementary Figure 6A**). Thus, the transmembrane domain of NRX and the short cytoplasmic domain are obligatory for formation of the ternary core-module, while the extended extracellular portion of the NRX protein is not able to promote stable core-module assembly.

We complemented these biochemical experiments with ‘molecular docking’ studies performed with AlphaFold2. The results predicted a 3D-structural arrangement (Methods) for the membrane-proximal regions of the ternary core-module where the transmembrane domains of NRX and PTPR are juxtaposed and surrounded by three of the four transmembrane segments of T178B (**Figure 2F**, left panel; **Supplementary Figure 6B**). In this framework, T178B is expected to closely link the two major classes of sCAMs, and to generate a macro-molecular complex with an extended surface for protein-protein interactions both in the membrane plane, as well in the aqueous phases on either side of the membrane (**Figure 2F**, right panel).

### Assembly in the ER and preferred localization of the core-module(s) to the plasma membrane

The structural arrangement of the ternary core-module and the association of its constituents with multiple partner proteins prompted the questions on where the co-assemblies of core-module(s) and partners occurs and what may happen upon targeted removal of the interlinking T178B protein.

For this purpose, we performed a series of experiments starting out with ‘organellar proteomics’ on 32 fractions derived from whole brain homogenates ((Schwenk et al., 2019), **Figure 3A**, upper panel) for probing the preferred sub-cellular localization of the individual core-module constituents. Analysis of the MS-derived molecular abundance values determined for all proteins in these fractions by t-SNE showed that all NRX paralogues and LAR-PTPRs are colocalized with the T178 proteins in a cluster delineated by established marker proteins of the plasma membrane ((Itzhak et al., 2016; Schwenk et al., 2019), **Figure 3A**). Similarly, a series of synaptic proteins such as AMPAR subunits, LRRTs, DLGs (also known as PSD proteins) or the family 2 voltage-gated calcium (Ca^2+^) channels (CAC1A (Cav2.1), CAC1B (Cav2.2), CAC1E (Cav2.3)) were found in this cluster, which itself appears clearly separated from other membrane compartments including ER, endosomes, Golgi apparatus or mitochondria (**Figure 3A**). This strongly preferred sub-cellular localization to the plasma membrane was further confirmed by meAP-MS experiments targeting the abundant post-synaptic LRRT4 protein that effectively retrieved the core-module(s) together with pore and auxiliary subunits of AMPARs ((Schwenk and Fakler, 2021; Schwenk et al., 2012), **Figure 3B**). Moreover, immunostaining of endogenous T178B detected via a knocked-in HA-epitope tag (Methods) independently confirmed localization of this tetraspanin to the synaptic compartment and co-localization with NRXs (**Supplementary Figure 7**).

**Figure 3.**
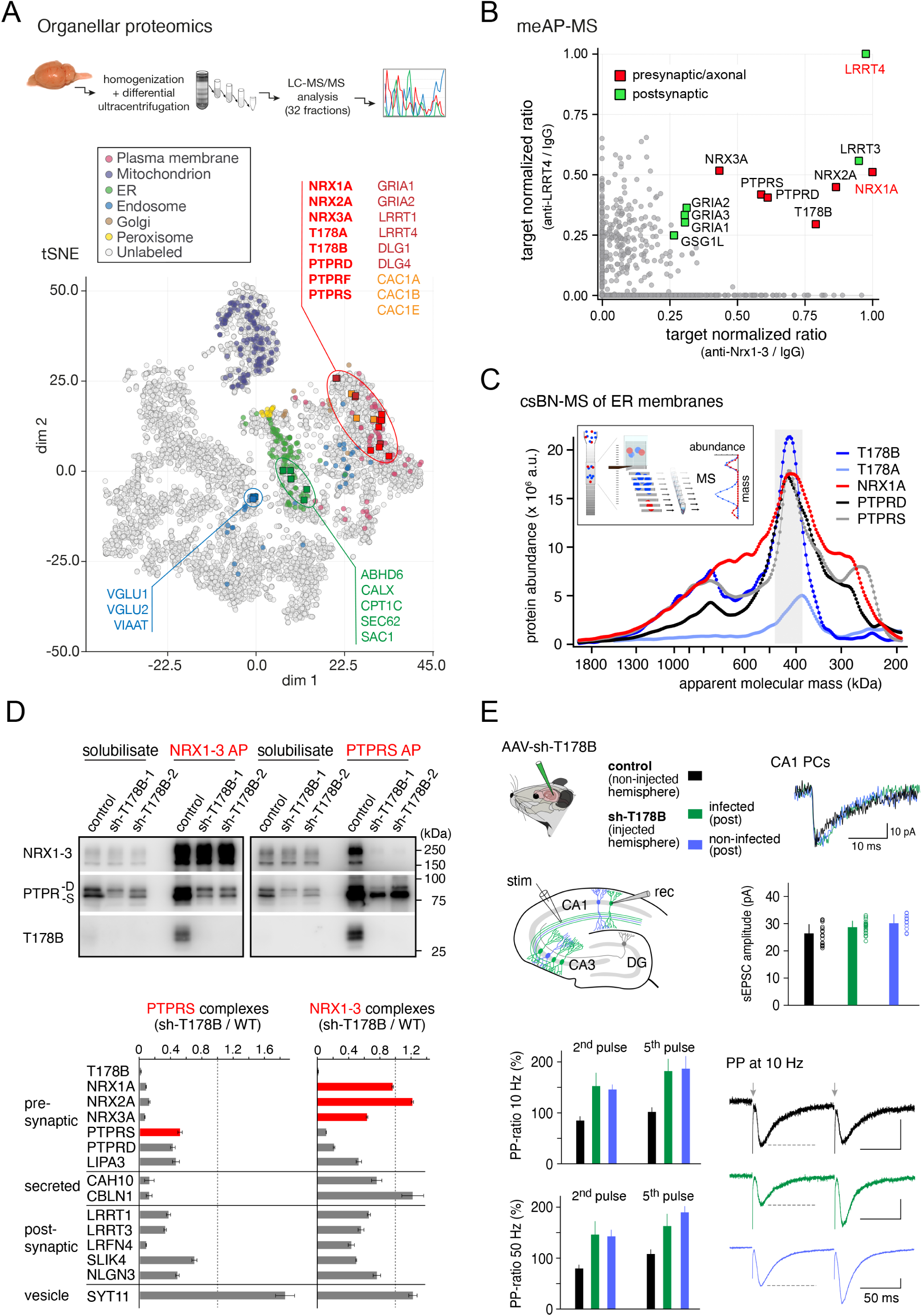
Distribution and biogenesis of core-module complexes and alteration of assembly and pre-synaptic functions upon knock-down of T178B. (A), Organellar proteomics of brain homogenates. Upper panel, key steps of the approach comprising fractionation of brain homogenates by density and differential sedimentation ultracentrifugation, quantitative MS of the 32 fractions and visualization of the relative protein abundance profiles by t-SNE. Lower panel, t-SNE plot of all individual MS-identified and quantified proteins (circles, total of 6738). Established markers for distinct organellar compartments (Itzhak et al., 2016) are color-coded as indicated. Circled areas indicate close co-localization of selected proteins in the plasma membrane (red), synaptic vesicles (blue) and endoplasmic reticulum (green). Note that all subunits of the ternary core-module show preferential localization at the synaptic plasma membrane. (B), tnR-plot of APs with ABs targeting post-synaptic LRRT4 and pre-synaptic NRX1-3 highlights common co-purification of PTPRs D, S and T178B and subunits of AMPA-type glutamate receptors. tnR-values were determined with IgGs as a reference. (C), Abundance-mass profiles obtained for the indicated proteins by csBN-MS in CL-47 solubilized ER-enriched membrane fractions from adult mouse brain. Inset: Scheme of csBN-MS approach used (Muller et al., 2019; Schulte et al., 2023). Note close co-migration of NRX1A, PTPRD,S and T178A, B indicating their co-assembly into complexes of variable apparent mass. Values of protein abundance indicate close to stoichiometric association (see text). (D), Alterations in PTPR and NRX complexes after knockdown of T178B. Upper panel, Input and eluates of APs with anti-PTPRS and anti-NRX1-3 APs from CL91-solubilized membrane fractions of cultured neurons transduced with two different AAV-sh-T178B viruses or control virus (WT) Western-probed with antibodies specific for NRX1-3, PTPRS,D or T178B. Lower panel, Abundance ratios determined for proteins specifically purified (and co-purified) in three anti-PTPRS and anti-NRX1-3 APs from WT and AAV-sh-T178B-2 transduced cultures by quantitative MS-analysis. Data are mean ± SD of three experiments, values for the AP targets are marked in red. (E), Alterations in pre-synaptic function induced by AAV-sh-T178B in hippocampal neurons of adult mice four weeks after stereotactic injection (inset, upper left). Upper right panel, Bar graph summarizing the amplitudes of AMPAR-mediated sEPSCs and representative EPSC traces measured in whole-cell recordings in CA1 pyramidal cells (CA1 PCs) in the un-injected hemisphere (control, WT, black) and in infected (green) and non-infected (blue) PCs of the injected hemisphere (inset, lower left). Data are mean ± SEM of 19 (WT), 13 (infected) and 12 (non-infected) PCs (p-values <0.01 between control and sh-injected hemisphere, Mann-Whitney U-test). Lower panel, Bar graphs summarizing paired-pulse (PP) ratios determined in CA1 PCs with stimulation frequencies of 10 and 50 Hz and representative AMPAR-mediated current traces recorded at 10 Hz, color-coding of expression manipulation as in the upper panel. Data are mean ± SEM of 11 (WT), 6 (infected) and 8 (non-infected) PCs (p-values <0.01 between control and sh-injected hemisphere, Mann-Whitney U-test). Note the marked increase in PP-ratio of AMPAR-currents triggered by sh-T178B infected pre-synapses in both infected and non-infected CA1 PCs.

The observed localization of the vast majority of the core-module constituents (NRXs, LAR-PTPRs and T178) to the plasma membrane does, however not exclude their co-assembly at earlier stages of membrane protein biogenesis. We, therefore, investigated their molecular appearance in ER-enriched membrane fractions (Schwenk et al., 2019) by complexome-profiling via csBN-MS, a technique that combines native gel electrophoresis with quantitative MS (**Figure 3C**, inset, (Kollewe et al., 2021; Muller et al., 2019; Schulte et al., 2023)). The ‘abundance-mass profiles’ obtained with this technique demonstrated effective co-assembly of NRX1A, T178B and PTPRS, D in the ER, most prominently reflected by the profile-peaks at an apparent mass of around 400 kDa (**Figure 3C**, shaded), comparable to the result obtained with the plasma membrane-enriched fractions before (**Figure 2A**). The abundance-peak of T178A at a slightly smaller mass (∼360 kDa) together with the broadening (‘shoulder’) of the peaks in the LAR-PTPRs and NRX1A profiles argues for preferred co-assembly of this tetraspanin with variants of the PTPRs and NRXs. Interestingly, the ER-profiles also showed formation of higher-mass assemblies (up to 1.2 MDa) indicating association of the core-module with further interactors. We explored the subunits of these assemblies by additional APs with anti-NRX1-3 and anti-PTPRS ABs and identified association with liprin-α2 (LIPA2), Cav2.2 (CAC1B), CAH10 and CBLN1 amongst other proteins (**Supplementary Table 3**). Thus, our csBN-MS data identified formation of the core-module during biogenesis in the ER and suggested its delivery to the plasma membrane as a pre-formed entity together with co-assembled partners.

### Re-organization of NRXs in the pre-synapse upon knock-down of T178B

Next, we explored the consequences of targeted knockdown of the NRX-PTPR-linking T178B protein by virally-driven expression of two different shRNAs (sh-T178B-1, -2) in both neuronal cultures and hippocampal neurons in mice.

In cultured neurons, AAV-delivered sh-T178B decreased the amount of the T178B protein present in (un-solubilized) membrane fractions by more than 95% as determined by quantitative MS (**Supplementary Figure 8**, **Supplementary Table 4**). More detailed analysis showed that this decrease was target-specific leaving the vast majority of proteins including the NRX paralogues in this fraction either unaltered or changed by ∼10% or less (**Supplementary Figure 8**, red lines). In contrast, the knockdown of T178B resulted in a reduction of both PTPRD and S by ∼30-40% (**Supplementary Figure 8**).

These changes in ‘expressed protein amounts’ translated into profound alterations in the composition of the core-module and its assemblies uncovered by anti-PTPRS and anti-NRX1-3 APs (**Figure 3D**). Thus, (1) the ternary complexes were largely abolished, leaving NRX and PTPRs either as separate proteins, or, as a minor portion of bi-molecular NRX-PTPR assemblies, (2) NRX lost 30-60% of its postsynaptic interactors such as LRRTs or NLGNs, as well as PTPR-linked presynaptic partners such as liprins (LIPA3) or proteins LRFN (leucine-rich repeat and fibronectin type-III domain containing), and (3) PTPRS was largely deprived of its NRX-recruited interactors including the CBLNs (**Figure 3D**).

The significance of the knockdown-induced changes was further investigated in the hippocampus of adult mice where the AAV-sh-T178B was delivered into the hippocampus (CA3 region, **Supplementary Figure 9**) of one hemisphere by stereotactic injection, while the other was left untouched for the use as wildtype control (WT, Methods). As illustrated in Figure 3E, whole-cell recordings performed in acute slices detected AMPAR-mediated excitatory post-synaptic currents (EPSCs) in response to spontaneous action potentials in pyramidal neurons of the CA1 region (CA1 PCs) in both hemispheres with similar time course and amplitude (**Figure 3E**, upper panel). Interestingly, EPSCs evoked by electrical stimulation of the Schaffer collaterals (**Figure 3E**, inset), the axons of CA3 PCs serving as presynaptic input to the CA1 PCs, uncovered marked differences between non-injected and virus-injected hemispheres. While control CA1 PCs responded with constant EPSCs in paired-pulse (PP) recordings (PP-ratio ∼100%) at stimulation frequencies of 10 and 50 Hz, PCs in the injected hemispheres displayed a marked stimulation-dependent increase in their EPSC amplitudes (PP-ratios of 150-200%; **Figure 3E**, lower panel). This increase in PP-ratio was induced by the knock-down of T178B in the pre-synaptic terminals, but was independent from knock-down in the post-synaptic neuron, pointing towards altered presynaptic function(s) in the absence of T178B (**Figure 3E**, lower panel, **Supplementary Figure 9**).

Together, these results on selective removal/reduction of T178B indicated the importance of this protein for formation of the ternary NRX-T178-PTPR complexes and demonstrated the significance of their structural integrity for normal synaptic transmission. Notwithstanding, they also showed that the remaining entities enabled operation of pre-synapses, albeit with characteristics that were different from normal presumably as a consequence of altered protein arrangements/assemblies.

This hypothesized re-organization of the core-module(s) upon knockdown of T178B was further pursued by analyzing the distribution of the NRXs in freeze-fracture replicas of the hippocampal CA1 region of WT and sh-T178B-injected mice by electron microscopy combined with immuno-gold-labelling with NRX1-3 ABs (Althof et al., 2015; Booker et al., 2020; Boudkkazi et al., 2023; Schwenk et al., 2019). Quantitative assessment of gold-particles showed that NRX1-3 exhibit a strong preference for (localization to) the active zones of both excitatory and inhibitory synapses identified by parallel labelling for the vesicular transporters of glutamate (VGLU1) and GABA (VIAAT), respectively (**Figure 4A, B, E**). Noteworthy, the gold-particles demonstrated tight embedding of the NRXs into the presynaptic protein network(s) reflected by the densely packed intra-membrane particles (IMPs), very similar to Cav2.1 (P/Q-type) channels (**Figure 4D**). Recapitulation of these experiments following virally-driven knockdown of T178B in the hippocampus largely abolished the preference of NRXs for the synaptic active zones, resulting in almost equal distribution of the NRX-particles between active zone (synaptic) and extra-synaptic localizations (**Figure 4C, E**).

**Figure 4.**
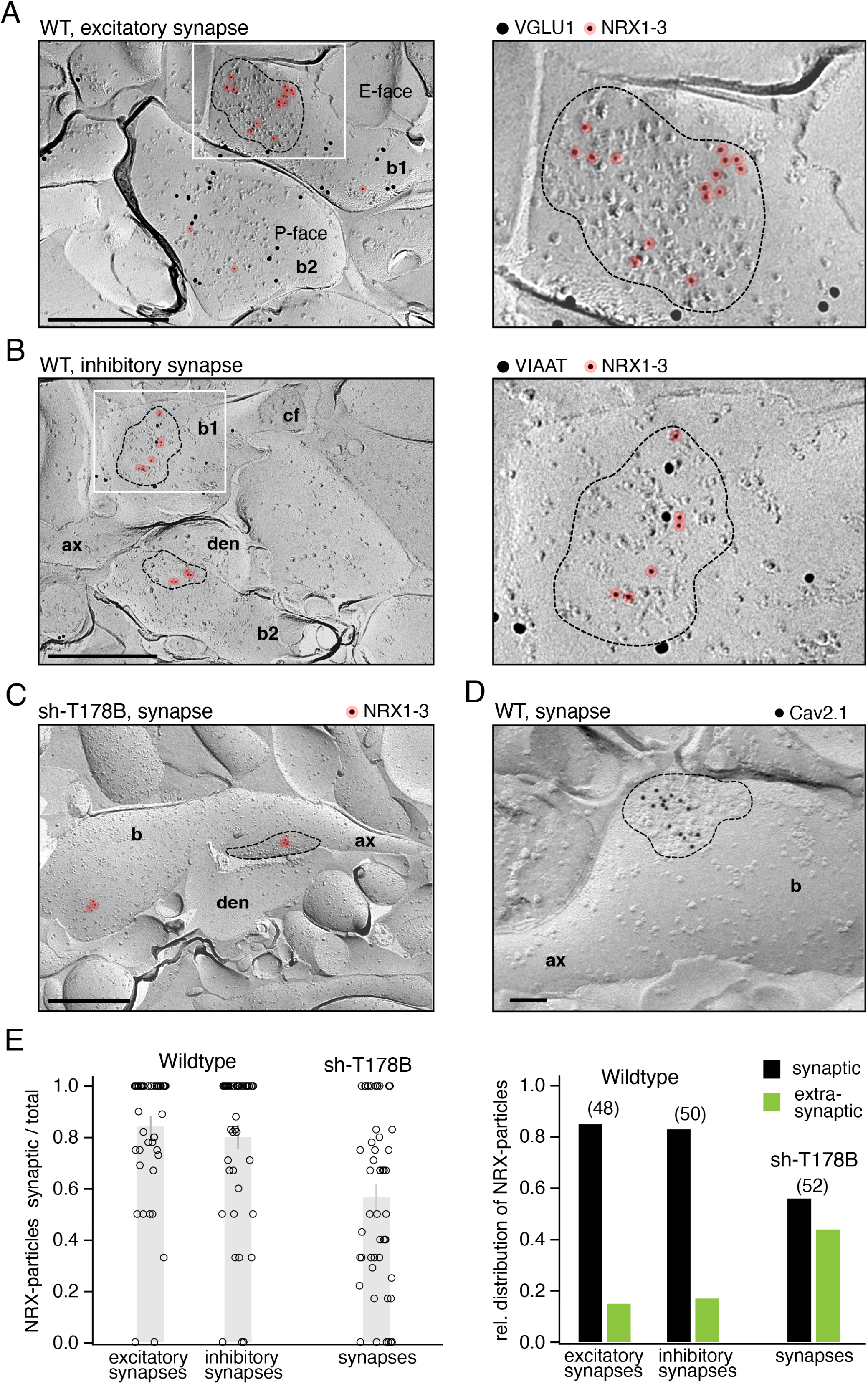
Localization of NRX1-3 to presynaptic active zones and re-distribution following knockdown of T178B. (A,B), Left panel, electron micrographs illustrating predominant localization of immuno-gold particles labelling NRX1-3 (6 nm particles, red overlay) to the active zone (delineated by broken line) and occasionally to the extra-synaptic membrane of excitatory and inhibitory synaptic boutons (b) in the hippocampal CA1 region of wildtype mice. Immuno-particles for the vesicular transporters of glutamate (VGLU1, 18 nm particles) and GABA (VIAAT, 18 nm particles) were used as markers for excitatory and inhibitory terminals, respectively. Right panel, framed area on the left at extended scale. In (B), NRX1-3 particles were additionally observed in the active zone of a VIAAT-negative putative excitatory bouton (b2). (C), Immuno-gold particles for NRX1-3 (12 nm, red overlay) localized to both the active zone (delineated by broken line, making synaptic contact with a postsynaptic dendrite (den)) and to an extra-synaptic membrane area of a presynaptic bouton. (D), Clustered organization of Cav2.1 channels (12 nm particles) in the active zone of a presynaptic bouton. (E), Quantitative assessment of the distribution of NRX1-3-particles to synaptic and extra-synaptic membranes of excitatory and inhibitory terminals in hippocampal CA1 regions of WT and sh-T178 injected mice. Data obtained from 48-52 boutons are illustrated individually (left panel, empty circles, grey bars indicate mean ± SD of the individual terminals) or as pooled data obtained in all terminals (right panel). Abbreviations: ax, preterminal axon; cf, cross-fractured face of bouton; P-face, protoplasmic face; E-face, exoplasmic face. Scale bars are 500 nm.

Together, these immuno-EM results demonstrated preferred localization of NRXs to the active zone of synaptic boutons as previously observed (Szoboszlay et al., 2017) and strongly suggested that this preference is due to co-assembly with T178B.

### Integration of the Neurexin-T178-PTPR module into pre- and trans-synaptic protein networks

To test whether the NRX-T178-PTPR core selectively engages in specific trans-synaptic interactions or whether it represents a more universal synapse-building block, we performed a series of reverse meAP-MS-experiments targeting an array of NRX-interactome constituents (**Figure 1**, **Table 1**). In total, 30 different constituents were selected as targets prioritized to cover (1) all three synaptic compartments, pre- and post-synaptic membrane, or the synaptic cleft and (2) the diverse physiological functions annotated in literature and public databases (**Figure 5**, upper panel, **Table 1**). Experimentally, all 64 reverse target meAPs (plus 27 control APs) were performed and rigorously evaluated as described before; in particular, specificity of co-purification (in APs with 1-4 ABs per target) was determined from MS-based tnR-values with target-KOs and/or pre-immunization IgGs as negative controls and an empirically-determined tnR of 0.25 (**Supplementary Figure 3A**) as a specificity threshold (Kocylowski et al., 2022b; Kollewe et al., 2022). The complete sets of proteins determined as specific interaction partners of a given target, together with all details related to the respective meAP experiments are summarized in Supplementary Table 4.

**Figure 5.**
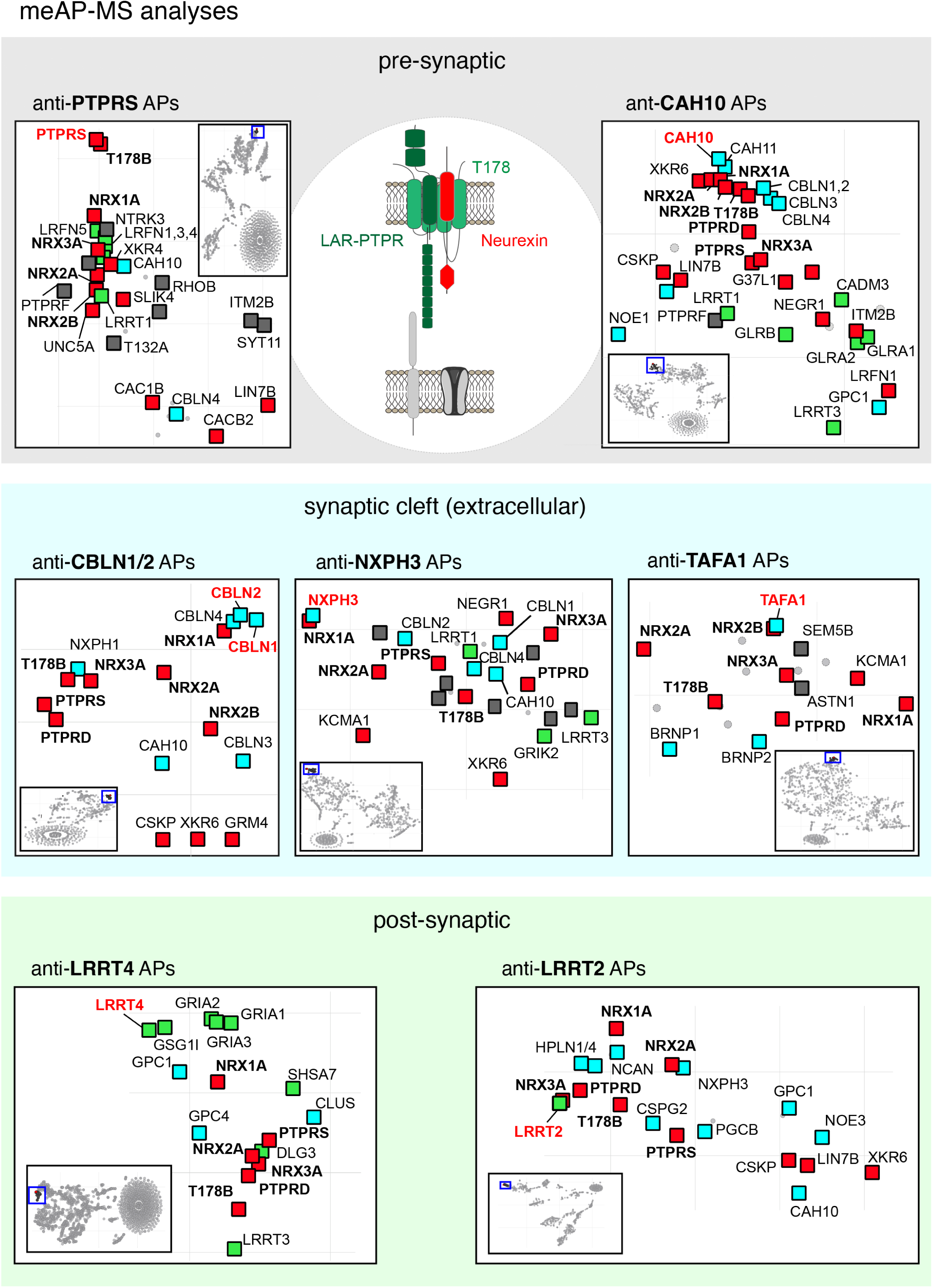
Comprehensive meAP-MS analyses uncover stable formation of trans-synaptic network(s) t-SNE plots of proteins resolved by their tnR-values in APs with multiple ABs specifically targeting the indicated NRX interactors in pre-synapse (PTPRS, CAH10), synaptic cleft (CBLN1/2, NXPH3, TAFA1) and post-synapse (LRRT4). For LRRT2 APs recombinant LRRT2 was used as a bait. Insets illustrate entire maps of the t-SNEs with all MS-detected proteins (for extension see **Supplementary Table 5**), enlarged sections (framed in blue) focus on the cluster of proteins in close proximity to the respective target protein. Note that PTPRD/S, T178B and NRXs appear as central elements in all clusters, while the set of proteins specifically and consistently co-purified with the respective target protein is variable. tnR values were determined with IgGs as controls. CBLNs were purified with anti-HA ABs from transgenic HA-Cbln1 and HA-Cbln2 mouse brains with wild-type brains serving as negative control.

Figure 5 illustrates the t-SNE plots of proteins positioned by their tnR-values obtained in multiple APs with six of the 30 selected targets (**Figure 5**, central inset): in the pre-synapse (PTPRS and CAH10, a protein covalently bound to NRX1-3 (Sterky et al., 2017)), in the synaptic cleft (CBLN1/2, neurexophilin 3 (NXPH3) and TAFA1) and in the post-synaptic membrane (LRRTs 2, 4). Similar to the anti-NRX1-3 APs (**Figure 1C**), these t-SNE plots of reverse APs readily delineated well-defined clusters of proteins identified as specific and consistent target-interactors by their close co-localization with the respective target (**Figure 5**, insets, blue frame(s)). This finding immediately indicated that all the NRX-associated target proteins were affinity-isolated as constituents of larger assemblies built on robust detergent-resistant protein-protein interactions. Remarkably, all these assemblies contained the molecularly diverse constituents of the NRX-T178-PTPR core-module (**Figure 5**) highlighting their common incorporation into the examined synaptic protein complexes. Beyond the ternary core-module, the t-SNE-delineated assemblies contained two groups of additional constituents. One group of proteins was shared among several of the isolated assemblies and thus likely forms an integral part of the central NRX-based building block(s), such as CBLNs, CAH10/11, or XKR6. The other group was uniquely detected with individual targets thus indicating their co-assembly with the NRX-based building block(s) in a more target-dependent manner. These selectively associated proteins represented a variety of distinct functional classes including proteins involved in cell adhesion, neuronal/axonal growth and development, voltage- and ligand-gated ion channels, G-protein coupled receptors (GPCRs), lipid-modulation, vesicle-fusion/processing or extracellular matrix (ECM). As an additional finding of major importance, the t-SNE plots showed that pre-synaptic targets (PTPRS and CAH10-NRXs) robustly co-purified integral post-synaptic membrane proteins (including LRRT1,3,4, and the AMPAR-subunits GRIA1-3, GSG1l, SHSA7, and NOE1,3 (Schwenk et al., 2012)), and vice versa. Accordingly, the NRX-T178-PTPR core-modules establish proteinaceous trans-synaptic bridges that are stable enough for effective quantitative affinity-isolation and that firmly link pre- and post-synaptic membrane proteins (**Figure 5**).

Together, these reverse meAPs strongly suggested that the NRX-T178-PTPR module(s) are the central hub elements of extended protein networks that expand within and across the synaptic membranes and that are built from a conserved core and a molecularly variable periphery.

### The synaptic connectome of the Neurexin-T178-PTPR core module

For more insight into the structure and complexity of these networks, we next explored their strings and (sub)-clusters by detailed analyses of the core module-associated interactomes (**Table 1**, **Supplementary Table 5**) and by using the quantitative data on protein abundances for automated cluster-analysis with the UMAP-algorithm (McInnes et al., 2018) (Methods, **Supplementary Figure 10**).

Figure 6 projects the resulting multi-protein network onto the synaptic topology and depicts its architecture built on the protein-protein interactions documented by specific and effective co-purifications between individual or groups of constituents (lines in **Figure 6**). Thus, the NRX-T178-PTPR module with its variable subunit composition forms the network core which, as an entity, promotes co-assembly with a multitude of constituents in all three sub-compartments of the synapse (**Figure 6**, **Supplementary Figure 10**). In the pre-synapse, the core-module robustly co-assembles with the key elements of the active zone (including UN13A, LIPA2/3, voltage-gated Ca^2+^ channels of the Cav2-family, RIM-binding proteins 1, 2 (RIMB1,2) and BK_Ca_-type K^+^ channels (KCMA1), **Supplementary Table 5**) and thus recruits the distinct components of the release-machinery into close spatial proximity (Emperador-Melero et al., 2024). Conversely, the core-module, alone or together with secreted constituents (CAH10, NXPH3), associates with excitatory and inhibitory transmitter-receptors or receptor complexes (AMPARs or kainate receptors, and GABA_A_-receptors (GABA_A_Rs) or glycine receptors) in the post-synaptic membrane thus juxtaposing them to the presynaptic release sites. Interestingly, network integration of the two major neurotransmitter-receptors, AMPARs and GABA_A_Rs, is profoundly different: While AMPARs are linked to the core-module through several different strings (NOE1-NRX-CAH10, GSG1l-LRRT4), GABA_A_Rs predominantly rely on the NRX-NLGN binding (**Figure 6**).

**Figure 6.**
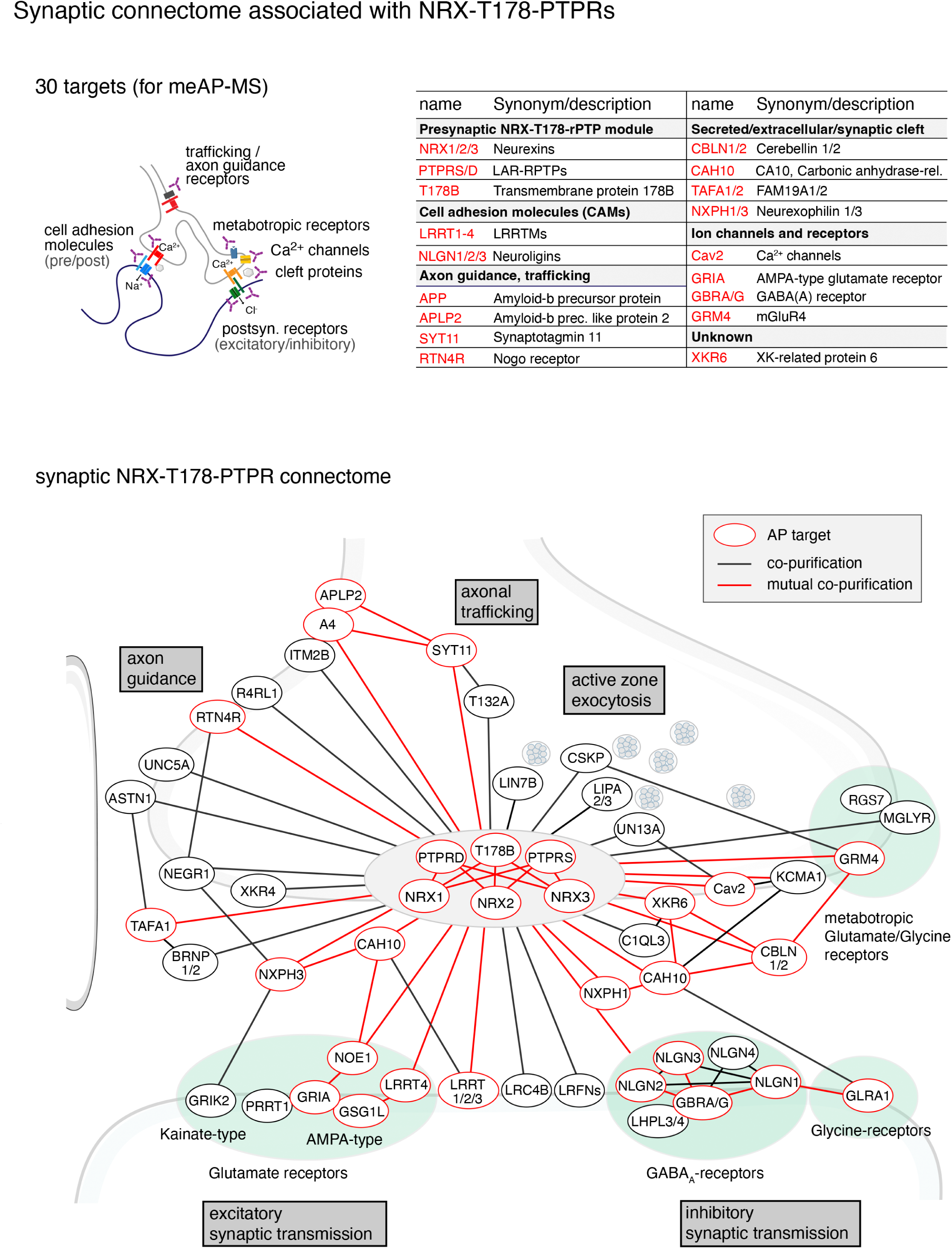
Architecture of NRX-associated protein networks projected onto the synaptic compartment. Graphical illustration of the architecture of NRX-T178-PTPR-associated networks built on robust and specific (co)-purifications in APs against the indicated 30 targets (upper panel) projected onto the synaptic compartment(s). Proteins which were specifically co-purified in individual APs with the respective target proteins (red circles) from brain membrane fractions were connected with lines in black, mutual co-purifications are indicated by red lines. Subunits/components of excitatory glutamate receptors, inhibitory GABA_A_- or glycine receptors and presynaptic GPCRs are highlighted by filled circles in green; overlapping circles indicate documented co-assembly of the respective proteins. Note the tight and multi-dented linkage/connection between the presynaptic NRX-T178-PTPR module and the post-synapse, as well as with the synaptic cleft. An interactive version of Figure 5 visualizing all ‘connectivities’ determined for any selected network constituent and providing additional details on AP-data and linkages to the UniProt/SwissProt database is available online (https://phys2.github.io/synaptic-connectome).

In addition to these network constituents operating in synaptic transmission, the NRX-T178-PTPR module co-assembles with a variety of constituents that may either serve ‘more specific’ functions (not present in any type of excitatory/inhibitory synapse) or currently lack functional annotations. These include the metabotropic glutamate receptor GRM4, and the metabotropic glycine receptor (MGLYR/GPR158)-RGS7 complex(Laboute et al., 2023), as well as several surface receptors previously implicated in neuronal development (and axonal pathfinding) such as RTN4R, R4RL1, ASTN1, NEGR1, the A4/APP-APLP2-ITM2B complex(Schwenk et al., 2016) or T132A, a presumed regulator of integrins and the integrin pathway (Li et al., 2022). Finally, the XKR proteins 4 and 6, either verified (XKR4) or possible (XKR6) lipid scramblases, should be mentioned based on the presumed significance of lipid signaling in (pre-)synaptic physiology (Neniskyte et al., 2023). The significance of these ‘more specific’ network constituents for synaptic organization and function remains to be resolved using the NRX-T178-PTPR module-instructed networks as a guide; for more detailed and ready accessibility, the network illustrated in **Figure 6** is, therefore, also provided as a an interactive figure version online (https://phys2.github.io/synaptic-connectome).

## Discussion/Conclusion

Individual CNS synapses contain an array of sCAMs but it remained enigmatic whether different classes of sCAMs have individual, additive contributions to synapse formation or whether they are tied into functional platforms. In this work, we uncovered that the two major classes of sCAMs predominantly assemble into a tightly-linked, universal pre-synaptic core-module for synapse formation, thus, providing a paradigm shift in the operation of synapse-organizing factors. Importantly, this NRX-T178-PTPR core-module acts as the central hub of solid trans-synaptic protein networks that recruit various classes of signaling entities in the pre-synapse and form solid high-affinity associations with postsynaptic receptors and adhesion systems.

### Molecular appearance of native synaptic protein networks - technical aspects of the proteomic analyses

For thorough and unbiased analysis of the molecular appearance of synaptic protein complexes in the adult rodent brain, we applied a largely refined proteomic workflow of meAP-MS that promotes comprehensiveness and specificity of the pursued interactomes (and interaction partners) by means of its key features (Kocylowski et al., 2022b; Kollewe et al., 2022; Schmidt et al., 2017; Schwenk et al., 2012; Schwenk et al., 2016): (1) use of solubilization conditions that preserve native higher molecular mass complexes of the target (with careful assessment of solubilization via BN-PAGE and MS-analysis (Kocylowski et al., 2022; Kollewe et al., 2022; Muller et al., 2010; Schulte et al., 2023; Schwenk et al., 2019; Schwenk et al., 2012; Schwenk et al., 2010), (2) application of multiple verified ABs with distinct epitopes on each target (**Figure 1**, **Supplementary Tables 1, 5**), (3) MS-based protein quantification with a linear range of up to four orders of magnitude and unbiased identification of all proteins, (4) the use of stringent negative controls (knockout input, target-unrelated ABs) and (5) consistency among ABs for determination of interactome constituents (visualized by t-SNE plots of tnR-values, **Supplementary Figure 3**, (Kocylowski et al., 2022)). Related to criteria (4) and (5), it must be noted that our meAP-MS approach aims at high reliability and a minimal number of false-positive interactors, deliberately operating at the cost of false negatives.

As an immediate outcome, this approach identified stoichiometric co-assembly of NRXs1-3 with all three LAR-type PTPRs and the previously uncharacterized tetraspanins T178A and T178B (**Figures 1-3**). In the ternary complexes, that form during early biogenesis in the ER (**Figure 3C**, see also below), the two tetraspanins act as structural ‘brackets’ stabilizing the otherwise rather loose bimolecular interactions between NRX and LAR-PTPRs (**Figure 2**). As a second main outcome, our approach uncovered the NRX-T178-PTPR module(s) as the central hub of trans-synaptic protein networks with exquisite stability and extension as defining features. Thus, the core-module recruited networks were resistant to detergent-treatment, appeared as large multi-component ‘entities’ (**Figures 1, 2, 4**) and could be effectively (large abundance values of the constituents) and consistently isolated from membrane preparations with multiple ABs targeting a variety of different constituents (visualized in t-SNE plots, **Figures 1, 4, 6**). Noteworthy, the isolated network entities uncovered solid trans-synaptic bondings between the presynaptic NRX-T178-PTPR module and several integral membrane proteins in the post-synapse. Finally, the architecture of the identified trans-synaptic synaptic network(s) was derived from all ‘entities’ isolated in extensive meAPs and were built on connectivities between individual constituents exclusively defined by specific, often mutual, co-purification(s) as a proxy for robust protein-protein interactions (**Figure 6**).

### Presynaptic protein networks – dependence of NRXs-localization on T178B

This embedding/integration of the NRX-T178-PTPR module into protein networks of the pre-synapse was immediately visualized by immuno-EM experiments on freeze fracture replicas of the hippocampal formation: The NRX proteins were found with strong preference (of > 80%) in the active zone areas in the midst of high densities of IMPs representing transmembrane proteins (**Figure 4**). This localization preference was independent of the type of synapse, although IMP densities are different for excitatory and inhibitory synapses. Most notably, the preference in active zone-localization strongly depended on the presence of T178B as demonstrated by the profound re-distribution of the NRXs following knockdown of T178B (**Figure 4C**, **E**). This re-distribution of the NRXs into areas with low IMP numbers is meaningful in two ways: First, it reflects the profound loss of NRX-interaction partners observed in meAP-MS experiments (**Figure 3D**), and, second, it provides a plausible explanation for the altered characteristics in synaptic transmission evidenced by the markedly increased paired-pulse ratios seen in knockdown animals but not in WT under our experimental conditions (**Figure 3E**).

In addition, the immuno-EM experiments, together with biochemistry/meAPs (**Figure 3D**) and patch-clamp recordings (**Figure 3E**) clearly demonstrated that removal of the T178-bracket modifies, but does not abolish synapse-development/formation, very similar to what has been reported for various individual or triple knockouts of NRXs or LAR-PTPRs (Anderson et al., 2015; Chen et al., 2017; Emperador-Melero et al., 2021; Horn et al., 2012; Missler et al., 2003; Takahashi and Craig, 2013). Rather, the core-module is of fundamental importance for the organization of pre-synaptic functions and trans-synaptic alignments (see below). In this respect, it will be important to further investigate the precise localization (and function) of the core-module constituents once the respective tools are in hand, in particular specifically and effectively labelling ABs for PTPRs and the two T178 isoforms, as well as the tagged variants of the latter that are currently in progress.

### Significance of the NRX-T178-PTPR module for synapse operation

The NRX-T178-PTPR module-associated synaptic network is made-up from an extensive number of constituents located in all three synaptic compartments. Both stability and extension of this network impact the operation of the synapses in several ways, beyond the significance of individual interactions.

Most importantly, the trans-synaptic network provides a solid ‘spatial organization platform’ where co-operating proteins are placed in close proximity and are thus enabled to functionally interact in a reliable and temporally defined manner. This is directly evident for the co-assembly of key active zone elements (Cav2, UN13A, LIPAs) with the NRX-T178-PTPR core in the pre-synaptic membrane and the core-module mediated recruitment of several neurotransmitter receptor complexes in the post-synaptic membrane. The resulting ‘nano-column’ arrangement (Biederer et al., 2017; Tang et al., 2016) juxtaposes release-site(s) and effectors of neurotransmitters and, thus, guarantees fast and reliable synaptic signal transduction. Remarkably, for AMPARs the trans-synaptic connectivity is realized by several molecularly distinct network strings (**Figure 6**) pointing to different types of synapses or different states of development and/or maturation. Similar synergism resulting from spatial proximity may be expected for other signaling/regulatory elements identified as co-assembled constituents of the NRX-T178-PTPR network, including the GPCRs GRM4 and MGLYR, the ion channel KCMA1 or the ‘synaptic organizers’ LRRTs or LRFNs (DeWit et al., 2013; Lie et al., 2018; Soler-Llavina et al., 2013; Um et al., 2016; Woo et al., 2009), but also the considerable number of proteins that have not yet been related to the two major cell adhesion molecules NRXs and LAR-PTPRs or that lack annotation of resolved primary functions (**Figure 6**, **Table 1**, **Supplementary Table 5**). This includes the putative scramblase XKR6, as well as proteins T132A, ASTN1, BRNP1, LRC4B or R4RL1.

### Significance of the NRX-T178-PTPR module for synapse assembly

The ternary NRX-T178-PTPR complex identified in this work uncovers an unexpected link between two major presynaptic hubs that were considered independent synaptic building blocks in the past (Gomez et al., 2021; Jang et al., 2017; Sudhof, 2017; Takahashi and Craig, 2013; Takahashi et al., 2012; Yuzaki, 2018). This has major consequences for interpreting the molecular logic of neuronal synapse assembly. First, the stoichiometric bundling of NRXs and PTPRs substantially expands the molecular interaction codes at the synapse by positioning two structurally distinct classes of adhesion cues into one ternary module. Second, the multivalent extracellular and intracellular interactions with this core module provide a mechanistic explanation for the extraordinary resilience of the synapse formation process in mammals to genetic perturbation. Notably, individual and even combined deletions of sCAM proteins examined in past studies only partially impair intracellular and extracellular protein interactions with the core-module. Moreover, we discovered that NRX1γ, a short NRX isoform not deleted in the previously reported ‘pan-Neurexin knock-out models’ (Sclip and Sudhof, 2023), is one of the most abundant neurexin isoforms in the mouse brain. Through the T178 tetraspanin link, this short protein is still sufficient to build NRX-T178-PTPR modules and nucleate pre- and trans-synaptic assemblies.

In aggregate, this work highlights unanticipated avenues for exploring the structural and functional underpinnings of trans-synaptic nanoscale organization and molecular recognition events at neuronal synapses. Moreover, the data provided here serve as a detailed roadmap for analyzing the role(s) of the network as an assembly platform (and the resulting spatial co-assembly/localization), as well as for studying the significance of the newly identified constituents with known and yet unknown function in synapse assembly, operation and dynamics.

## Acknowledgements

We thank the Fakler and Scheiffele lab members for helpful discussions and I. Schaber for excellent technical assistance. B.F. appreciates the interactions with T. Südhof and his efforts leading to initiation of this project.

## Author Contributions

J.S., U.S. and B.F. conceived the project, S.T., J.S., P.H., M.K.K. and U.S. did all the experiments related to protein biochemistry and proteomic analyses, S.B., A.B. and S.T. performed electrophysiology, reconstruction of neurons and immune-cytochemistry, N.S and A.K. carried out immuno-EM analysis, E.M. and A.H. generated T178B-HA mice and did immunostainings, J.S., A.H. and J.vdB. performed data analysis and set up the interactive Figure 6, D.K., F.S. D.S., M.Y. and P.S. generated and characterized KO/transgenic animals, B.F. evaluated data and wrote the manuscript together with J.S. and P.S., supported by all authors.

## Data and availability

Mass spectrometry data from AP-MS analyses, membrane proteomics and organellar proteomics have been deposited in PRIDE (PXD054017 and 10.6019/PXD054017). The meAP-MS data reconstituting the network illustrated in Figure 5 are provided as an interactive version (https://phys2.github.io/synaptic-connectome). All other data reported in this work are available from corresponding authors upon request.

## Methods

### Generation of knockout/knockin animals

HA-NRX1 knock-in mice (Sterky et al., 2017; Trotter et al., 2019), NRX3 knock-out (generation see **Supplementary Figure 5**), NRX3 AS5 HA/HA (Hauser et al., 2022), HA-Nlgn1 knock-in mice (Nozawa et al., 2018) and HA-Cbln1 (Nozawa et al., 2022) were generated as described previously. NRX3 knock-outs (**Supplementary Figure 5**) were generated by inserting a gene-trap after the first exon common to both α- and β-isoforms, using homology-arms PCR-amplified from C57BL/6 genomic BAC clones. The targeting vector, containing an SDA (self-deletion anchor)-flanked neomycin resistance cassette, and a DTA (diphtheria toxin fragment A)-cassette for negative selection, was electroporated into C57BL/6 ES cells (Cyagen). G418-resistant clones were isolated, verified by PCR and Southern Blot and injected into C57Bl/6 albino embryos (Cyagen). Following germline transmission and deletion of the neomycin cassette, heterozygotes were intercrossed to generate homozygous Nrxn3^gt/gt^ offspring and wildtype littermates. HA-Cbln2 knock-in mice were generated following the approach described in (Nozawa et al., 2022). Briefly, superovulated B6D2F1 female mice (8 weeks old) were mated with B6D2F1 males and zygotes were collected from the oviducts at embryonic day 0.5 and incubated with SpCas9 protein (250ng/µl; PNA Bio, CA, USA), Cbln2-crRNA (5pmol/µl, 5’-CGUAAGGGCUCAGAACGACAGUUUUAGAGCUAUGCUGUUUUG-3’), trans-activating RNA (5pmol/µl; Fasmac, Kanagawa, Japan) and 131-bases single-strand donor DNA oligo(500ng/µl; 5’-TGGCCCTGCTGTTGCTGCTGCTG-CCCGCCTGCTGCCCCGTAAGGGCTCAGTACCCATACGACGTGCCAGACTACGCT AACGACACGGAGCCCATCGTGCTAGAGGGCAAGTGCCTGGTAGTGTGCGATTCC-3’; Integrated DNA Technologies, IA, USA) at 37℃ for 15 min. Electroporation was performed using platinum block electrodes (150 mA, 2 pulses, 1msecON+100msecOFF, GE-101, BEX, Tokyo, Japan) connected to CUY21EDIT II (BEX). Zygotes were incubated to the two-cell stage and transferred into oviducts of presudo-pregnant ICR females. To investigate CRISPR/Cas9-mediated mutation in the Cbln2 gene, the genomic DNA was prepared from mouse tales and the genomic regions flanking the gRNA target were amplified by PCR using specific primers: Fwd (5’-ACTTGTTCCCCTTCTACCTGCC-3’) and Rev (5’-GTACGGTTGCTCA-TCTCTGACG-3’). Knockouts for CA10 (*Car10*) were generated by breeding *Car11^em1Sud^ Car10^em1Sud^/J* mice (Jackson Laboratories #031998) to C57Bl/6 and subsequently intercrossing *Car10^em1Sud^* heterozygotes to obtain homozygous offspring and wildtype littermates.

T178B-HA knock-in mice were generated by electroporation as described previously (Nozawa et al., 2018)with minor modifications. C57BL/6J fertilized eggs were incubated with Alt-RTMHiFi SpCas9 Nuclease V3 (250 ng/ml; Integrated DNA Technologies, IA, USA), crRNA (5 pmol/μl, 5’-CCU AAA UCC AGA CCA GAG AAG UUU UAG AGC UAU GCU GUU UUG -3′), trans-activating RNA (5 pmol/µl; Fasmac, Kanagawa, Japan ) and a single-stranded oligodeoxynucleotide (500 ng/μl; 5’-CCT CTG TCC AAC CTG TCC CGA GGA CCA ACT GCC CTA AAT CCA GAC CAG AGG GAT CCT ACC CAT ACG ACG TGC CAG ACT ACG CTA ATG GGA CAG TGT GCT AAA AGA CAA ACA AAC CCA TAC CTA TAT ATA TAT A -3’; Integrated DNA Technologies), which harbored a sequence encoding a hemagglutinin (HA) epitope tag and a BamHI site (GGATCC) flanked by homologous arms corresponding to exon 4 of *Tmem18B*, at 37 °C for 15 min before electroporation. Electroporation was performed using platinum block electrodes (100 mA, [3 msec On + 97 msec Off] for 6 pulses; GE-101, BEX, Tokyo, Japan) connected to CUY21EDIT II (BEX). Zygotes were incubated at 37 °C in 5% CO_2_ until they reached the two-cell stage and transferred into oviducts of pseudo-pregnant ICR females. Mice were housed with a 12:12 h light-dark cycle with food and water available ad libitum. The sex for immunohistochemistry was not distinguished.

### Molecular Biology

The cDNAs used were all verified by sequencing. T178B carrying a Myc-Flag Tag at the C-terminus was obtained from Origene (Klon RR216273) and subcloned into pcDNA3.1. PTPRS (GenBank: BC083188.1) was modified with a V5-tag at the C-terminus. HA-tagged NRX1A subcloned into pCAG was obtained from Addgene (#58622). HA-tagged constructs for expression of NRX3A and NRX3A AS5 were as described in (Hauser et al., 2022).

### Biochemistry and Cell Biology

#### Membrane preparation from mouse brain

Whole brains were dissected from adult C57BL/6 WT mice (Charles River) or genetically modified mice. Brains were cut into pieces and homogenized in 10 mM Tris/HCl buffer (pH 7.5) supplemented with 320 mM sucrose, 1.5 mM MgCl_2,_ 1 mM EGTA, 1 mM Iodoacetamide and protease inhibitors (Aprotinin, Leupeptin, Pepstatin A, PMSF) using a loose pestle in a 15 ml dounce tissue grinder. Supernatants of centrifugations (4 min, 1,000xg) were collected in tubes for ultracentrifugation. Homogenization procedure was repeated with the remaining tissue remnants in order to increase the yield. Combined supernatants were ultracentrifuged (200,000 x g, for 20 min). The resulting pellet were homogenized with dounce tissue grinder (tight pestle) in hypotonic lysis buffer (10 mM Tris/HCl (pH 7,4) and protease inhibitors) and incubated on ice for 20 minutes. After ultracentrifugation (200,000 x g, for 30 min) membrane pellets were resuspended in 8 ml of 0.5 M sucrose / 20 mM Tris/HCl (pH 7.4) and homogenized. Suspension were filled in thin layer ultracentrifugation tubes and underlaid with 12 ml 0.5 M sucrose / 20 mM Tris/HCl (pH 7.4) and 18 ml 1.3 M sucrose / 20 mM Tris/HCl (pH 7.4) and subjected to ultracentrifugation (180,000 x g, for 30 min, SW-32, Beckman). The membrane phase between the two sucrose layers was harvested and diluted with 20 mM Tris/HCl (pH 7.4) to wash out sucrose. Ultracentrifugation (200,000 x g, for 30 min) was used to pellet the membranes which were then resuspended in a small volume of 20 mM Tris/HCl (pH 7.4) to reach concentrations about 10 mg/ml. Protein concentration were determined by Bradford assay and samples were aliquoted and shock-frozen in liquid nitrogen. For proteomic inspection of protein abundances in membrane fraction from adult mouse brains. Equal amounts of proteins obtained from 4 individually prepared wild-type brains were denatured by adding 10 µl Laemmli-buffer, separated on SDS-PAGE and silver stained.

Bands were cut at 45 kDa and 140 kDa and the three gel pieces were subjected to tryptic digest for nano-LC MS/MS (**Figure 1B, D**).

#### Preparation of ER-enriched mouse brain fraction

The isolation of ER-enriched membrane fractions from mouse brain was essentially done as described in (Schwenk et al., 2019). For complexome profiling (csBN-MS) was adjusted as follows. Whole brains of C57BL/6 WT mice were dissected and homogenised in 0.25 M SHE buffer (10 mM HEPES (pH 7.5), 1 mM EDTA, 2 mM PMSF, protease inhibitors see above) first using a loose pestle in a 15 mL dounce tissue grinder and then processed in a cell cracker (8.002 mm ball, 2 passages, 2 rounds per turn). Homogenates were centrifuged at 600 x g for 5 min. The supernatant was collected and the pellet was washed with homogenization buffer. Combined supernatants were centrifuged again (600 x g for 10 min). The resulting supernatant was then subjected to ultracentrifugation (200,000 x g, for 30 min). The pellet (P200) was homogenized in hypotonic buffer (5 mM HEPES (pH 7.5) containing 1 mM PMSF and protease inhibitors), left to incubate for 1 hour on ice with agitation and subsequently centrifuged at 12,000 x g, for 15 min. The supernatant was collected and placed on top of a two layer density gradient (8.3% (0.25 M) and 23% buffered sucrose). After ultracentrifugation (135,000 x g for 1 h) the interface between 8,3% and 23% sucrose layers was harvested, diluted in 20 mM Tris/HCl (pH 7.5), and pelleted (200,000 x g, for 25 min). The ER-enriched membrane vesicles were resuspended in 20 mM Tris/HCl (pH 7.5). Protein concentrations were determined by Bradford assay.

#### Covalent coupling of antibodies

For APs, selected ABs were covalently coupled to magnetic Dynabeads ™ Protein A or Dynabeads ™ Protein G. Selected ABs were diluted in PBST (1 μg of AB with 10 μl filtered PBST) and incubated with Dynabeads™ Protein A or G, for 1 h (4 μl Dynabeads per μg of coupled AB). Subsequently, ABs coupled to beads were crosslinked for 30 min by addition of 20 mM Dimethyl pimelimidate dihydrochloride in pre-filtered 100 mM sodium borate, 0.05% Tween (pH 9). Reactions were blocked by adding 200 mM ethanolamine, 0.05% Tween (pH 8). Non-crosslinked ABs were washed off by incubation for 2 x 5 minutes with 100 mM Glycine in 0.05% Tween (pH 3). After 3 x washing steps with PBST coupled ABs were used for APs or stored for short periods at 4°C.

#### Antibody based affinity-purifications

Protein complexes in plasma membrane and ER enriched fractions from mouse brains were solubilised with ComplexioLyte 91 (CL-91) or ComplexioLyte 47a (CL-47, Logopharm GmbH) supplemented with 1 mM CaCl_2_ (and protease inhibitors (Aprotinin, Leupeptin, Pepstatin A, PMSF)). After clearing by ultracentrifugation (10 min, 125,000 x g) solubilisates (1-1.5 mg) were incubated for 2 hours with 10-15 µg ABs pre-coupled to protein A Dynabeads. APs and respective control experiments used to calculate the specificity of interactions (see tnR-ratio calculation below), were performed with identical settings. Target APs were replicated by using different batches of membrane fractions generated from WT mouse brains (as described above). AP experiments with genetically modified brain tissue as source material (NLGN1-HA, CBLN1-HA, CBLN2-HA, Nrx1-HA, NRX3-KO) were done in parallel with controls obtained from age-matched WT littermates. Details of the experimental series are described in the Figure legends and are given in Tables S1-4. The following antibodies were used for APs: *anti-Nrx1-3* (Millipore, #ABN161-I), *anti-Nrx1-3* (AB generation by LifeTein, #RB2987, rabbit, epitope: mouse NRX3A aa 1523-1571); *anti-Nrx1-3* (Synaptic Systems, #175003), *anti-Nrx1-3* (AB generation by AbFrontier, #3-4-rb1, rabbit, epitope: mouse NRX3A aa 1471-1491), *anti-Nrx1* (Synaptic Systems, #175103), *anti-Nrx1* (RnD Systems, #AF4524), *anti-Nrx2* (AB generation by LifeTein, #RB4227, rabbit, epitope: mouse NRX2A aa 266-282), *anti-Nrx2* (AB generation by LifeTein, #RB4226, rabbit, epitope: mouse NRX2A aa 1459-1476), *anti-Nrx2* (AB generation by LifeTein, rabbit, #RB4225, epitope: mouse NRX2A aa 1459-1476), *anti-Nrx2* (Abcam, #ab34245), *anti-Nrx3* (AB generation by AbFrontier, rabbit, #3-1-rb1, epitope: mouse NRX3A aa 1349-1368), *anti-Nrx3* (Synaptic Systems, #175303), *anti-Nrx3* (AB generation by AbFrontier, rabbit, #3-2-rb1, epitope: mouse NRX3A aa 1377-1397), *anti-Nrx3* (RnD Systems, #AF5269), *anti-Nrx3* (AB generation by AbFrontier, rabbit, #3-2-rb2, epitope: mouse NRX3A aa 1377-1397), anti-T178B (AB generation by LifeTein, rabbit, #RB4235, epitope: mouse T178B aa 273-294), anti-T178B (AB generation by AmsBio, rabbit, #RB3967, epitope: mouse T178B aa 273-294), anti-HA (Roche, #11867423001), anti-HA (Thermo Fisher, #26183), anti-PTPRS (RnD Systems, #AF3430), anti-PTPRS (MediMabs, #MM-0020-P), anti-PTPRS (Invitrogen, #PA5-47487), anti-CAH10 (RnD Systems, #MAB2189), anti-CAH10 (raised against recombinant CAH10 ((Montoliu-Gaya et al., 2021), rabbit, #Hadlai), anti-CAH10 (raised against recombinant CAH10 (Montoliu-Gaya et al., 2021), rabbit, #Habeba), anti-CAC1A (FrontierInstitute, #GP-Af810), anti-CAC1E (Alomone, #ACC-006), anti-TAFA1 (RnD Systems, #AF5154), anti-TAFA2 (RnD Systems, #AF4179), anti-NXPH1 (RnD Systems, #AF4208), anti-NXPH3 (RnD Systems, #AF5098), anti-LRRT4 (Neuromab, #75-261), anti-LRRT4 (AbFrontier, rabbit, #C1, raised against mouse LRRT4 aa 548-568), anti-LRRT4 (RnD Systems, #AF5377), anti-A4 (Abgent, #ABIN1741750), anti-A4 (Sigma, #A8717), anti-GBRA1 (Synaptic Systems, #224203), anti-GBRG2 (Synaptic Systems, #224003), anti-RTN4R (RnD Systems, #AF1440), anti-LRRT1 (MerckMillipore, #ABN642), anti-LRRT2 (NeuroMab, #75-364), anti-GRM4 (Abcam, #ab53088), anti-GRM4 (Abcam, #ab184302), anti-GRM4 (Bioworld, #BS1144), anti-NLGN2 (Synaptic Systems, #129511), anti-NLGN3 (NeuroMab, #75-158), anti-Syt11 (Synaptic Systems, #270003), anti-XKR6 (AB generation by AmsBio, rabbit, #RB3961, epitope: mouse XKR6 aa 85-102), anti-XKR6 (AB generation by AmsBio, rabbit, #RB3962, epitope: mouse XKR6 aa 85-102), anti-XKR6 (Signalway Antibody, #42864), IgG (Millipore, #12-370), rabbit pre-immune-sera (LifeTein). Similarly, 15 µg recombinant LRRT2 proteins (RnD Systems, #5589-LR-050) were used as affinity matrix for protein complex isolations after pre-coupling to protein A Dynabeads.

#### 2D Blue Native/SDS-PAGE analysis of complexes

Protein complexes in mouse brain plasma membrane-enriched fractions (Kollewe et al., 2022; Schwenk et al., 2019; Schwenk et al., 2016) were solubilized with CL-47 (1 mg membrane protein / 1 ml CL-47 supplemented with 1 mM PMSF, 1 mM Iodoacetamide and a mixture of protease inhibitors (Leupeptin, Pepstatin A, Aprotinin). After clearing by ultracentrifugation (125,000xg, 10 min.) solubilisates were supplemented with 0.05% Coomassie® Brilliant Blue G-250 dye (Carl Roth) and 8% Glycerol. Protein complexes were separated on a 3-15% (w/v) acrylamide linear gradient gel (format 1.6 x 140 x 80 mm), made with 50 mM BisTris (pH 7,5), 50 mM α-capronicacid and 0,01% GDN (Anatrace, #GDN101). Solubilisates (1 ml) were loaded on 3 cm pockets and separated by electrophoresis (180 Volt, 32 mA, 45 min, 600 V, 6 hours) with 15 mM BisTris / 50 mM Tricine / 0.02% Coomassie G250 as cathode and 15 mM BisTris (pH 7.4) as anode buffer. Gel lanes were excised, cut in strips of 0.8 cm, incubated at 37°C for 20 min in 0.1% SDS (pH 8.8), 100 mM DTT and horizontally placed on top of 10% SDS-PAGE gels. After electrophoretic separation and transfer on a PVDF membrane proteins were stained with Ruby Blot stain to visualize abundant protein complexes, which were used as internal molecular weight standard for estimation of the apparent molecular sizes in the first dimension. PVDF membranes were cut horizontally at 50 kDa and 125 kDa. The low molecular weight section was immunodecorated with anti-T178B (LifeTein, #RB4235), the 50-125 kDa section with anti-PTPR (Neuromab, #75-194) and the high-molecular area with anti-NRX1-3 (LifeTein, #RB2987). Antibodies were diluted 1:1000 in PBST + 3% BSA. Anti-HRP-conjugated secondary ABs (Abcam) in combination with ECL Prime (GE Healthcare) were used for visualization.

#### Enzymatic removal of heparan sulfates (HS)

Impact of HS on NRX-interactions was assessed by APs after pre-treatment with a heparinase cocktail. For this purpose, plasma-membrane enriched fractions from mouse brain were incubated for 1 h with a mix of Heparinases (2 Units/ml mix of Heparinase I (Sigma-Aldrich, H2519), Heparinase II (Sigma-Aldrich, H6512) and Heparinase III (Sigma-Aldrich, H8891)) diluted in reaction buffer (20 mM Tris/HCl (pH 7.4), 100 mM NaCl, 2 mM CaCl_2_ and a mix of protease inhibitors (Aprotinin, Leupeptin, Pepstatin A). As control, membranes processed in parallel without additions of enzymes were used. After enzymatic treatment, membranes were pelleted by ultracentrifugation (100,000 x g for 10 min, 4°C) and supernatants were removed. Subsequently membranes were solubilized with CL-91 and processed as described above for APs with a mixture of anti-Nrx1-3 antibodies (anti-Nrx1, #ABN161, Millipore; anti-Nrx1, Synaptic Systems, #175103) and subjected to MS-analysis.

#### Complex reconstitution in tsA cells

tsA201 cells were used for transient co-expression of V5-tagged PTPRS, T178B-Myc-FLAG, NRX1A-HA and NRX3A-HA. Cells were cultivated in high glucose Dulbecco’s Minimal Eagles Medium (DMEM), supplemented with 0.45% glucose, 10% (v/v) FCS, 1% (v/v) Penicillin/Streptomycin and 1% (v/v) HEPES. Two days after PEI (Polysciences)-mediated transfection cells were washed with PBS and released by cell scrapers in 20 mM Tris/HCl (pH 7,4), 150 mM NaCl, 1 mM EDTA, 1 mM PMSF and protease inhibitors. Cells were centrifuged (150,000xg, 10 min) and lysed in 20 mM Tris/HCl (pH 7,4) by ultrasonication (5s-pulses, output 50%). Membrane vesicles were pelleted by ultracentrifugation (150,000xg, 10 min) and resuspended to homogeneity in 20 mM Tris/HCl (pH 7.4). Protein concentrations of membrane suspensions were determined by Bradford assay and aliquots shock-frozen in liquid nitrogen and stored at -80°C. Experiments were performed in parallel. For each AP, 0.5 mg membrane fractions were incubated for 30 min in CL-91 (supplemented with 1 mM PMSF, 1 mM Iodoacetamide). After clearing by ultracentrifugation (125,000xg, 10 min), solubilisates were incubated for 2 hours with 3 µg of coupled ABs (V5, BioRad, #MCA1360) or 40 µl anti-FLAG affinity resin (Millipore, A2220). Affinity-matrices were briefly washed twice with detergent buffer before elution of bound proteins in 10 µl 1xSDS-buffer without DTT. Aliquots of solubilisates and eluates were run on 10% or 12% SDS-PAGE and proteins transferred onto PVDF membranes. Immunodecorations were done with anti-HA (Thermo Scientific, #26183), anti-V5 (BioRad, #MCA1360) and anti-FLAG (Thermo Scientific, #MA1-91878) antibodies diluted 1:1000 in PBST + 3% BSA. Anti-HRP-conjugated secondary ABs (Abcam) in combination with ECL Prime (GE Healthcare) were used for visualization.

### Mass Spectrometry

#### LC-MS/MS analysis

Vacuum-dried peptides were dissolved in 13 µl or 20 µl of 0.5% (v/v) trifluoroacetic acid. Appropriate amounts were loaded onto trap columns (C18 PepMap100, 5 μm particles, Thermo Fisher Scientific GmbH, Dreieich, Germany) with 0.05% trifluoroacetic acid and separated on C18 reversed phase columns (SilicaTip emitters, 75 μm i.d., 8 μm tip, New Objective, Inc, Littleton, USA, manually packed 11 to 12 cm (Orbitrap Elite mass spectrometer) or 21 to 22 cm (QExactive HF-X mass spectrometer) with ReproSil-Pur ODS-3, 3 μm particles, Dr. A. Maisch HPLC GmbH, Ammerbuch-Entringen, Germany; flow rate: 300 nL/min) using UltiMate 3000 RSLCnano HPLC systems (Thermo Fisher Scientific GmbH, Dreieich, Germany). Gradients were built with eluent _ˈ_A_ˈ_ (0.5% (v/v) acetic acid in water) and eluent _ˈ_B_ˈ_ (0.5% (v/v) acetic acid in 80% (v/v) acetonitrile / 20% (v/v) water): 5 min 3% _ˈ_B_ˈ_, 60 min from 3% _ˈ_B_ˈ_ to 30% _ˈ_B_ˈ_, 15 min from 30% _ˈ_B_ˈ_ to 99% _ˈ_B_ˈ_, 5 min 99% _ˈ_B_ˈ_, 5 min from 99% _ˈ_B_ˈ_ to 3% _ˈ_B_ˈ_, 15 min 3% _ˈ_B_ˈ_ (Orbitrap Elite mass spectrometer) or up to 5 min 3% _ˈ_B_ˈ_, 120 min from 3% _ˈ_B_ˈ_ to 30% _ˈ_B_ˈ_, 20 min from 30% _ˈ_B_ˈ_ to 40% _ˈ_B_ˈ_, 10 min from 40% _ˈ_B_ˈ_ to 50% _ˈ_B_ˈ_, 5 min from 50% _ˈ_B_ˈ_ to 99% _ˈ_B_ˈ_, 5 min 99% _ˈ_B_ˈ_, 5 min from 99% _ˈ_B_ˈ_ to 3% _ˈ_B_ˈ_, 10 min 3% _ˈ_B_ˈ_ (QExactive HF-X mass spectrometer). Eluting peptides were electro-sprayed at 2.3 kV (positive polarity) via Nanospray Flex ion sources into an Orbitrap Elite mass spectrometer (CID fragmentation of the 10 most abundant at least doubly charged new precursors per scan cycle) or into a QExactive HF-X mass spectrometer (HCD fragmentation of the 25 most abundant doubly, triply, or quadruply charged new precursors per scan cycle; Figs. 1B, 3B) (all Thermo Fisher Scientific GmbH, Dreieich, Germany) and analyzed with the following major settings: scan range 370 to 1,700 m/z, full MS resolution 240,000, dd-MS2 resolution _ˈ_normal_ˈ_ (ion trap) or 15,000, respectively, maximum dd-MS2 injection time 200 ms or 100 ms, respectively, intensity threshold 2,000 or 40,000, respectively, dynamic exclusion 30 s or 60 s, respectively, isolation width 1.0 m/z. LC-MS/MS RAW files were converted into peak lists (Mascot generic format, mgf) with ProteoWizard msConvert (https://proteowizard.sourceforge.io/). All peak lists were searched twice with Mascot Server 2.6.2 (Matrix Science Ltd, London, UK) against a database containing all mouse, rat, and human entries of the UniProtKB/Swiss-Prot database. Initially broad mass tolerances were used. Based on the search results peak lists were linear shift mass recalibrated using in-house developed software and searched again with narrow mass tolerances for high-resolution peaks (peptide mass tolerance ± 5 ppm; fragment mass tolerance 0.8 Da for Orbitrap Elite peak lists and ± 20 mmu for Q Exactive HF-X peak lists). One missed trypsin cleavage and common variable modifications were accepted. Default significance threshold (p < 0.05) and an expect value cut-off of 0.5 were used for displaying search results.

#### MS quantification of proteins

Proteins were evaluated by a label-free quantification procedure (Kocylowski et al., 2022). MaxQuant v1.6.3 (http://www.maxquant.org) was used for mass calibration and to extract from FT full scans peptide signal intensities (peak volumes, PVs). The elution times of peptide PVs were aligned (pairwise, Loess regression) and assigned to peptides by matching m/z and elution times acquired by MS/MS identification (tolerances 2-3 ppm / ± 1 min) as described (Bildl et al., 2012). The obtained PV tables (protein-specific peptide signal intensities in all runs) were then further processed to eliminate the influence of PV outliers, false assignments and gaps by exploring the consistency of PV relations within proteins (i.e. protein-specific PV ratios between and within runs). Orthogonal combinations of these relations provided ‘expected PV values’ (EPVs) for each interconnected PV value that served as a measure of accuracy and weighting factor. In addition, time- and run-dependent detectability thresholds were estimated for each cell in the PV table based on the distribution of measured PVs (3rd percentile within a 3 min elution time window). Qualified PV data in each protein PV matrix was then aggregated to obtain global protein references termed ‘protein reference ridges’ (i.e. vectors that represent the maximum protein coverage of MS/MS-identified and quantified peptides with their ionization efficiencies). In a final step, proteins were quantified by weighted fitting of their measured peptide PVs to their respective reference ridge. In case no (consistent) peptide PVs were identified, an apparent protein detection limit was determined using the detectability thresholds of the three best ionizing peptides fitted to the protein reference ridge. Abundance_norm_spec values (as a measure of molecular abundance) were calculated as described (Bildl et al., 2012) from the fitted protein ridges. This procedure is comparable to MaxQuant LFQ ((Cox et al., 2014); all data is fitted without weighting), but quantification is more robust and accurate, in particular for sparse PV data.

#### Evaluation of proteomic results

Determination of specificity in APs was based on target-normalized abundance ratios (tnRs) of proteins in target APs versus control APs (calculated as described in (Kocylowski et al., 2022a)) together with information on molecular abundance and detection thresholds of proteins. These information were collectively inspected using the BELKI software suite (https://github.com/phys2/belki). For large data sets tnR-values were visualized by t-distributed stochastic neighbour embedding (t-SNE, with perplexity of 50, **Figures 1C, 5**); comparison of two APs was illustrated as 2D-scatter plot of tnR-values (Figure 2B). Specific co-purification was defined by two criteria: First, tnR values of ≥ 0.25 were set as specificity-threshold (Kocylowski et al., 2022a), unless high-quality ABs or stringent negative controls (target knock-out) allowed reduce of the threshold to 0.2. Second, consistency (of specific co-purification) among APs was applied (**Table 1**, **Supplementary Tables 2-5**, **Figure 6**), i.e. co-purification of a given protein was dubbed specific, if its tnR-value(s) exceeded the specificity-threshold in APs with more than two (or the majority of) target ABs or in replicate experiments. Details about the experiments (ABs, controls), tnR-pairs and specificity-thresholds for each experimental series are indicated in **Supplementary Tables 2-5**. For evaluation of NRX isoform expression in membrane fractions from adult WT mouse brains, proteins were first separated on SDS-PAGE and sections covering molecular weight ranges from 0-45 kDa, 45-140 kDa and >140 kDa were cut and individually analyzed by MS (**Supplementary Figure 1**). Signals obtained in high molecular weight sections were assigned to NRX1/2/3 alpha. Sequences unique for beta isoforms were exclusively found between 45-140 kDa and, therefore, the respective fractions were used to determine abundances of NRX1/2/3 beta. Signals obtained in low molecular weight fractions matched with NRX1γ. Abundance_norm_ values were used to calculate relative abundances (**Figure 1B**).

For evaluation of whole membrane fractions from wild-type mouse brains or neuronal cultures, total protein amounts were equilibrated to correct for systematic deviations due to technical variations (e.g. sample loading, in-gel digest etc.); normalization factors were smaller than 1.3. Mean values of abundance_norm_spec values determined in n=4 individual mouse brain membrane fractions were calculated and plotted relative to mean abundance_norm_spec values obtained in eluates of four anti-NRX1-3 APs (2 experiments with anti-NRX1-3 (Millipore, #ABN161-I) and (LifeTein, #RB2987) respectively) (**Figure 1D**). For direct comparison, all four AP data sets were first normalized to target values.

Similarly, abundance_norm_spec values were used to judge the impact of GPI-modification in NRX3A on protein-protein interactions (**Figure 2C**), to reveal changes on NRX1-3 complexes by enzymatic removal of HS-moieties (Figure 2D), and to monitor alterations in NRX1-3/PTPRS complex formation after sh-RNA mediated knockdown of T178B (**Figure 3D**). Abundance ratios providing average fluctuations in association with specific target proteins. Experiments were done in triplicates and values are given as mean ± SEM.

Organellar proteomic analysis used the data published in (Schwenk et al., 2019) re-processed as follows. MS/MS data was searched against the UniProtKB/Swiss-Prot MOUSE ReferenceProteome database with the common modifications described above. Quantitative evaluation of the proteins identified with at least three specific peptides was carried out as detailed above. Relative abundance profiles of proteins together with information from Itzhak et al. (Itzhak et al., 2016) on general subcellular markers were visualized in BELKI (t-stochastic neighborhood embedding (t-SNE) with perplexity of 40).

For UMAP-analysis of the NRX-T178-PTPR associated networks (**Supplementary Figure 10**), abundance_norm_Spec values of 172 proteins, identified as specific interactors in target APs (30 targets outlined in **Figure 6** (except of GRIA) and replicate experiments, indicated in **Supplementary Tables 2-5**), were used to uncover data structures pointing towards clusters or preferred assemblies. The UMAP (Sainburg et al., 2021) method was used to reduce the dimensionality of the normalized data, leveraging the software provided by McInnes *et al*. (https://doi.org/10.21105/joss.00861), which is based in the programming language Python (Van Rossum and Drake Jr, 1995). The Euclidean distance was chosen as the metric for clustering the data. The parameter constraining the size of the local neighborhood of a data instance (n_neighbors) was 10. Additionally, the Python library pandas (<u>10.5281/zenodo.3509134</u>) was used for data handling. Visualization of reduced data was done using the Bokeh library (http://www.bokeh.pydata.org.)

#### ER complexome profiling

ER-enriched membrane fractions (Schwenk et al., 2019) were prepared from freshly isolated mouse brains and gently solubilized with CL-47 (salt replaced by 750 mM aminocaproic acid). After ultracentrifugation, 1 mg of solubilized protein was concentrated by ultracentrifugation into a 20%/50% sucrose cushion supplied with 0.125% Coomassie G250 Blue. The sample was then subjected to high-resolution cryo-slicing blue native gel mass spectrometry (csBN-MS) as described recently (Schulte et al., 2023). Briefly, the BN gel (1-18%) lane was sliced into 359 samples (0.3mm intervals), each digested with trypsin, and the obtained peptides (dissolved in 0.5% (v/v) trifluoroacetic acid) were measured on a QExactive mass spectrometer coupled to an UltiMate 3000 RSLCnano HPLC system (Thermo Scientific, Germany). After absorption on a C18 PepMap100 precolumn (300 µm i.d. × 5 mm; particle size 5 µm; 0.05% (v/v) trifluoroacetic acid; 5 min 20 µL/min), peptides were eluted with an aqueous-organic gradient (eluent A: 0.5% (v/v) acetic acid; eluent B: 0.5% (v/v) acetic acid in 80% (v/v) acetonitrile) as follows: 5 min 3% B, 120 min from 3% B to 30% B, 20 min from 30% B to 50% B, 10 min from 50% B to 99% B, 5 min 99% B, 5 min from 99% B to 3% B, 10 min 3% B (flow rate 300 nL/min). Separation column was a SilicaTip™ emitter (i.d. 75 µm; tip 8 µm; New Objective, USA) packed manually (23 cm) with ReproSil-Pur 120 ODS-3 (C18; particle size 3 µm; Dr. Maisch HPLC, Germany) from which eluting peptides were directly electrosprayed (2.3 kV; transfer capillary temperature 300°C) in positive ion mode. MS acquisition parameters were: maximum MS/MS injection time = 400 ms; dynamic exclusion duration = 60 s; minimum signal/intensity threshold = 40,000 (counts), top 15 precursors fragmented; isolation width = 1.4 m/z. Mass spectrometric raw data was processed as described above: mass offset-corrected peak lists were searched with 5 ppm mass tolerance against all mouse, rat, and human entries of the UniProtKB/Swiss-Prot database (release 20181205). Acetyl (Protein N-term), Carbamidomethyl (C), Gln->pyro-Glu (N-term Q), Glu->pyro-Glu (N-term E), Oxidation (M), Phospho (S, T, Y), and Propionamide (C) were chosen as variable modifications (fragment mass tolerance: 20 mmu), maximum one missed tryptic cleavage allowed; expect value cut-off for peptide assignment was set to 0.5. Proteins either representing exogenous contaminations (e.g., keratins, trypsin, IgG chains) or identified by only one specific peptide were not considered. Ion intensity information (peak volumes, PVs) extracted by MaxQuant and assigned to peptides after retention time-alignment of runs (RT tolerance 1 min, m/z tolerance 5 ppm) was used to compute protein abundance profiles as detailed in (Schulte et al., 2023). Slice numbers were converted to apparent molecular mass by fitting of log(MW) of marker complexes versus their observed peak maxima in abundance-mass profiles using a sigmoidal function. Abundance-mass profiles were finally smoothed using a gaussian filter (width set to 1.4).

### Neuronal culture and sh-mediated knockdown

Brains were dissected from P18 rat embryos. Hippocampi and cortices of both hemispheres were dissected and transferred into HBSS buffer. Tissue pieces were trypsinized and briefly washed with HBSS buffer after stopping the enzymatic treatment by adding FCS. Cells were dissociated by trituration with a fire polished Pasteur pipettes. Vital cells were plated at high density (500,000 cells/well) for biochemical experiments or medium density (50,000 cells/well) for immunofluorescence experiments on poly-D-Lysine coated wells. For immunostainings cells were cultivated on coated glass coverslips. Neurobasal medium conditioned by cultivated glia cells was added at the start of the culture and freshly every week. Growing of mitotic cells was blocked by supplementing the medium with 5 µM AraC.

Knockdown of T178B by sh-T178B was induced by adding AAV DJ8 particles carrying sequences for H1 promoter-regulated transcription of sh-T178B (sh#1: GCCAGGAAGCACAGGGACA; sh#2: ACTTTGAGCTATCACGTTA) and GFP, whose expression is controlled by an ubiquitin promoter. Control viruses were driving sole GFP expression. Successful infection and vitality of cultured neurons were controlled regularly; cultures were maintained 2.5 up to 3 weeks to ensure synapse maturation and a quantitative elimination of T178B. Neurons were then used either for immunostainings or biochemical experiments. For the latter, neurons were released by cell scrapers and homogenized in 10 mM HEPES, 320 mM sucrose (pH 7.4) and protease inhibitors by up and down pipetting. After ultracentrifugation (125,000xg, 20 min.) the cell pellets were homogenized in 20 mM Tris/HCl (pH 7.4) and protein concentrations were determined by Bradford. Equal amounts of proteins (control, sh-treated neurons) were denatured by adding 10 µl Laemmli-buffer, separated on SDS-PAGE and silver stained. Bands were cut in low and high molecular weight pieces and subjected to tryptic digest for nano-LC MS/MS (**Supplementary Figure 8**, **Supplementary Table 4**). For analysis (**Figure 3D**) of protein assemblies, neurons were solubilized in CL-91 buffer and NRX and PTPRS complexes were affinity-isolated and processed for MS-analysis as described above. In addition, small aliquots of solubilisates and eluates were run on 10% SDS-PAGE and proteins transferred onto PVDF membranes. Blots were cut at 50 and 125 kDa and then separately immunodecorated with anti-T178B (LifeTein, #RB4235), anti-PTPR (Neuromab, #75-194) and anti-NRX1 (Millipore, ABN161-I). Antibodies were diluted 1:1000 in PBST + 3% BSA. Anti-HRP-conjugated secondary antibodies (Abcam) in combination with ECL Prime (GE Healthcare) were used for visualization.

### Electron microscopy

Immuno-gold labelling of SDS-digested freeze-fracture replicas (SDS-FRL) was carried out as previously described (Booker et al., 2020). Briefly, coronal hippocampal sections (120 µm) were cut from brains of two adult male WT mice and an adult male sh-T178B injected mouse which had been transcardially perfused with 0.9% NaCl followed by a fixative containing 1% paraformaldehyde (PFA) and 15% saturated picric acid in 0.1 M phosphate buffer (PB; pH 7.4). Slices were cryoprotected with 30% glycerol in 0.1 M PB overnight (O/N) at 4°C, then blocks containing the strata oriens, pyramidale and radiatum of CA1 microdissected and frozen with high pressure freezer (HPM 100, Leica, Austria). Frozen samples were fractured at -140°C, then coated by deposition of carbon (5 nm), platinum/carbon (2 nm) and carbon (18 nm) in a freeze-fracture replica machine (ACE 900, Leica). Replicas were subsequently digested for 18 hours (hrs) at 80°C in a solution containing 2.5% SDS and 20% sucrose diluted in 15 mM Tris buffer (TB; pH 8.3). Replicas were washed in a washing buffer (0.05% bovine serum albumin (BSA, Roth, Germany) and 0.1% Tween-20, in 50 mM Tris-buffered saline /TBS/), blocked in a solution containing 5% BSA and 0.1% Tween-20 made up in TBS for 1 hr at room temperature (RT). Subsequently, replicas, obtained from WT animals, were incubated with one of the following mixtures of primary ABs: (i) vesicular glutamate transporter 1 (VGLU1, goat (Go), 0.5 µg/ml, Frontier Institute, Hokkaido; (Kusch et al., 2018)) and NRX1-3 (LifeTein, #RB2987, rabbit (Rb), 3 μg/ml, see above), (ii) vesicular GABA transporter (VIAAT, Guinea pig (Gp), 4.5 µg/ml, Frontier Institute, Hokkaido; (Althof et al., 2015)) and NRX1-3 made up in 50 mM TBS containing 1% BSA and 0.1% Tween-20 (antibody solution) O/N at 15°C. After washing replicas were reacted with a mixture of 6 nm gold-coupled donkey anti-rabbit IgG and either (i) 18 nm gold-coupled donkey anti-goat IgG or (ii) 18 nm gold-coupled anti-guinea pig IgG secondary antibodies (1:30; Jackson ImmunoResearch Europe, Cambridgeshire, UK) diluted in antibody solution O/N at 15°C. NRX1-3 single labeling was performed on replicas obtained from T178B KO mouse. Antibody reaction was completed with 12 nm gold-coupled donkey anti-rabbit IgG secondary antibody (1:30; Jackson ImmunoResearch). All ABs used targeted intracellular epitopes and, therefore, result in labeling of the protoplasmic face (P-face), but not exoplasmic face (E-face). Replicas were subsequently rinsed in TBS, ultrapure water and then mounted on 100-mesh grids.

For quantitative analysis, VGLU1- and VIAAT-positive, as well as single NRX1-3 labeled boutons were imaged at 10,000x magnification under an electron microscope (Jeol JEM 2100 Plus, Jeol, Japan). To differentiate between synaptic and extrasynaptic membrane segments of boutons, active zones of the terminals were identified based on the typical morphology consisting of concave shape of the P-face and the higher density of intramembrane particles (IMPs) compared to the surrounding extrasynaptic membrane areas (Martin-Belmonte et al., 2025). Only active zones with clear morphological features were selected for the analysis. Finally, the relative distribution of NRX1-3 particles was calculated in synaptic and extrasynaptic areas of the boutons.

### Immunofluorescence

Mice were transcardially perfused with 4% paraformaldehyde (PFA) in 0.1 M phosphate buffer. Sagittal sections (50 μm thickness) were prepared using a microslicer (DTK-1000, Dosaka EM, Kyoto, Japan). For staining of synaptic proteins, sections were treated with 1 mg/ml pepsin in 0.2 N HCl/PBS for 5 min at 37 °C. Sections were incubated with 10% normal donkey serum, followed by primary ABs overnight, and then Alexa 405, 488, 647(Molecular probe, Thermo Fisher), and Alexa 555 (Abcam, Cambridge, UK)-conjugated species-specific secondary ABs for 2 h. Fluorescence images were taken by confocal microscopy (FV-100, Evident, Tokyo, Japan). The following ABs were used: anti-HA (rat, Roche, # 11867423001, RRID: AB_390918), anti-pan Neurexin (rabbit, Nittobo Medical Co., Ltd, # MSFR104640, RRID: AB_2571817), anti-PSD95 (guinea pig, Nittobo Medical Co., Ltd, MSFR105190, RRID: AB_2571612), anti-guinea pig IgG-Alexa Fluor 488 (donkey, Jackson ImmunoResearch Labs, # 706-545-148, RRID: AB_2340472), anti-rat IgG-Alexa Fluor 555 (donkey, Thermo Fisher Scientific, # A78945, RRID: AB_2910652), anti-rabbit IgG-Cy5 (donkey, Jackson ImmunoResearch Labs, # 711-175-152, RRID: AB_2340607).

### Transcript expression pattern

Transcript levels of individual genes in defined cell-types were calculated using the online software DropViz (Saunders et al., 2018). For this purpose, all subclusters assigned to a particular cell-type, according to the common name function of DropViz, were combined to a meta-group using the cluster comparison feature of DropViz with default parameter settings. The RNA count of a selected gene per 100,000 unique molecular identifiers in the meta-group was then given as target sum per 100k in a downloadable table containing differentially expressed genes.

### Electrophysiology

#### In Vivo Stereotactic Injection

The distinct AAVs were injected into C57BL/6 mice 20–23 days after birth (P20-P23). Animals were anesthetized by injection of a ketamine/dorbene mixture and mounted in a Kopf stereotaxic frame (Tujunga). Virus-containing solution (0.5–1 μl) was injected at a single site into the hippocampus (targeting the CA3 region) by means of a UMP3 controller (WPI, Sarasota) and a nanofil syringe/needle (WPI, Sarasota). Following surgery, pups recovered rapidly by antagonist injection and were returned to their home cage. Recordings were performed 4-5 weeks following virus injection. Animal procedures were in accordance with national and institutional guidelines and approved by the Animal Care Committee Freiburg according to the Tierschutzgesetz (AZ G-21/40).

#### Slice preparation

Transverse 300-μm-thick hippocampal slices were cut from brains of 1-to 2-month-old WT mice, as described (Boudkkazi et al., 2014). Hippocampal slices were cut in ice-cold, sucrose-containing physiological saline using a commercial vibratome (VT1200S, Leica Microsystems). Slices were incubated at 35°C, transferred into a recording chamber, and super-fused with physiological saline at room temperature.

#### Patch-clamp recordings in brain slices

Patch pipettes were pulled from borosilicate glass (Hilgenberg; outer diameter, 2 mm; wall thickness, 0.5 mm for somatic recordings). When filled with internal solution, they had resistances of ∼2–4 MΩ postsynaptic pipettes. Patch pipettes were positioned using Kleindiek micromanipulator (Kleindiek Nanotechnik, Reutlingen). A Multiclamp 700B amplifier (Molecular Devices, Sunnydale) was used for recordings. Pipette capacitance of both electrodes was compensated to 70%-90%. Voltage and current signals were filtered at 10 kHz with the built-in low-pass Bessel filter and digitized at 20 kHz using a Digidata 1440A (Molecular Devices, Sunnydale). pClamp10 software (Molecular Devices, Sunnydale) was used for stimulation and data acquisition. Holding potential of CA1 pyramidal neuronal was -70 mV (to record AMPAR-mediated EPSCs). Synaptic responses were evoked by stimulation at 0.1 Hz with a monopolar glass electrode filled with ACSF in the stratum radiatum of CA1 for CA1 pyramidal neurons. To ensure stable recording, membrane holding current, input resistance, and pipette series resistance were monitored throughout the recording. Statistical significance of differences was assessed by non-parametric Mann-Whitney U-test (as indicated in the legend of Fig. 3).

#### Solutions

For dissection and storage of slices, a sucrose-containing physiological saline containing 87 mM NaCl, 25 mM NaHCO_3_, 25 mM D-glucose, 75 mM sucrose, 2.5 mM KCl, 1.25 mM NaH_2_PO_4_, 0.5 mM CaCl_2_, and 7 mM MgCl_2_ was used. Slices were superfused with physiological extracellular solution that contained 125 mM NaCl, 25 mM NaHCO_3_, 2.5 mM KCl, 1.25 mM NaH_2_PO_4_, 1 mM MgCl_2_, 2 mM CaCl_2_, and 25 mM glucose (equilibrated with a 95% O_2_/5% CO_2_ gas mixture). Pipettes were filled with a K-methylsulfonate intracellular solution containing 120 mM KMeHSO_3_, 20 mM KCl, 2 mM MgCl_2_, 2 mM Na_2_ATP, 10 mM HEPES, and 0.1 mM EGTA. Membrane potentials are given without correction for liquid junction potentials. Values given throughout the manuscript indicate mean ± SEM or SD. Significance of differences was assessed by a nonparametric Mann-Whitney test.

## Supplemental Material

### Supplementary Figures

**Supplementary Figure 1.**
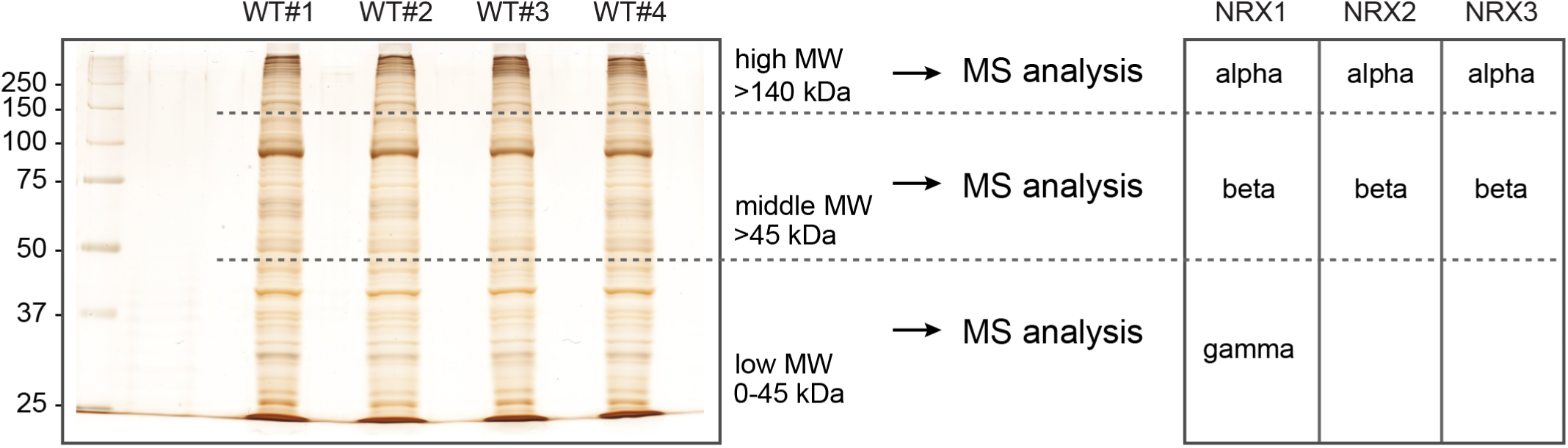
Protein abundances of NRX isoforms. Relative abundance of NRX isoforms (alpha, beta, gamma) was determined by quantitative MS in un-solubilized membrane fractions prepared from adult mouse brains. Proteins were separated by denaturing SDS-PAGE, the gel was cut at 45 and 140 kDa and protein abundance was determined in individual sections. All peptides matching the NRX1γ sequence in the 0-45 kDa sections were used to calculate the abundance of this NRX proteoform. Similarly, protein abundances of NRX1β and NRX1α were determined from the middle and high MW sections, respectively.

**Supplementary Figure 2.**
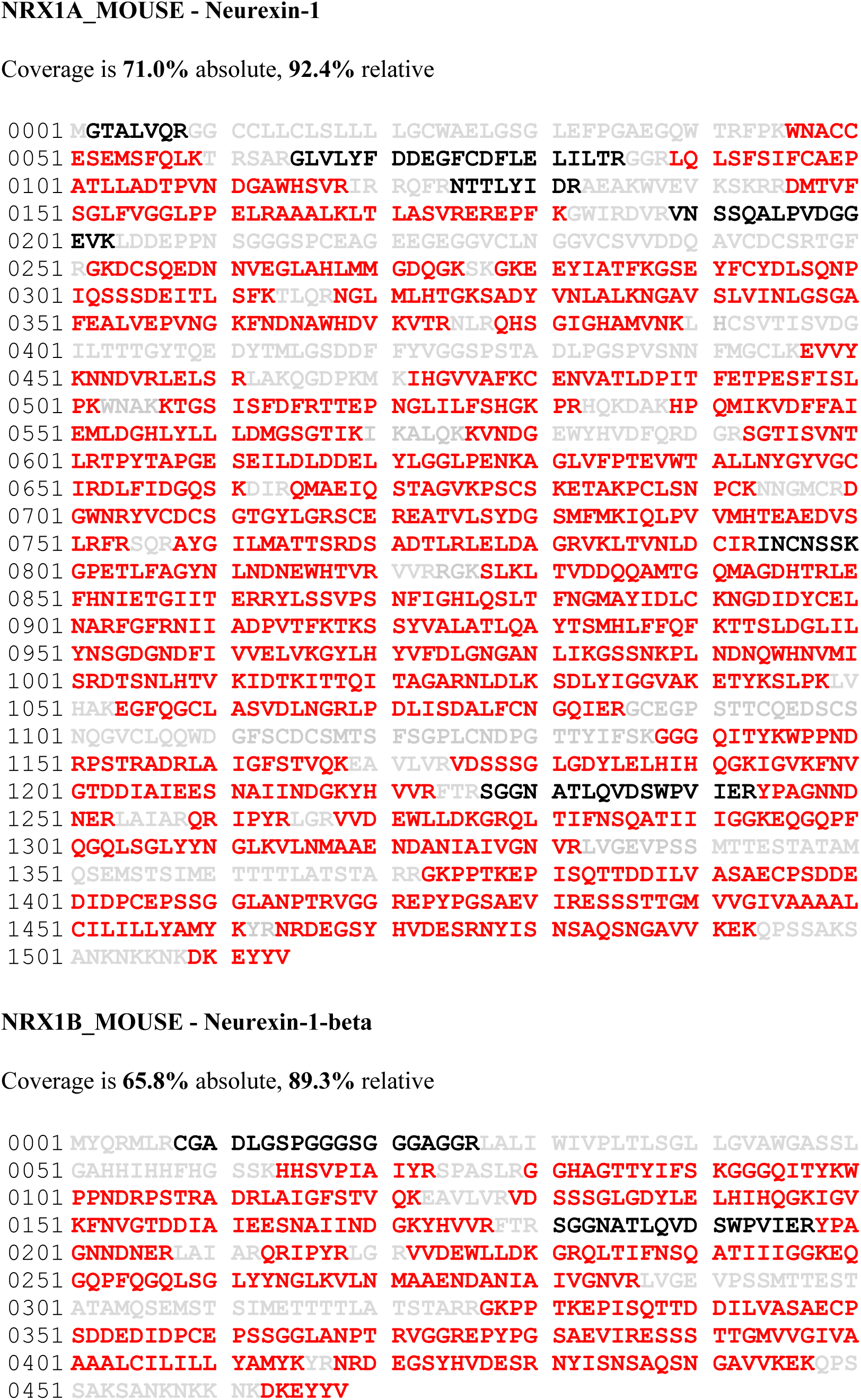
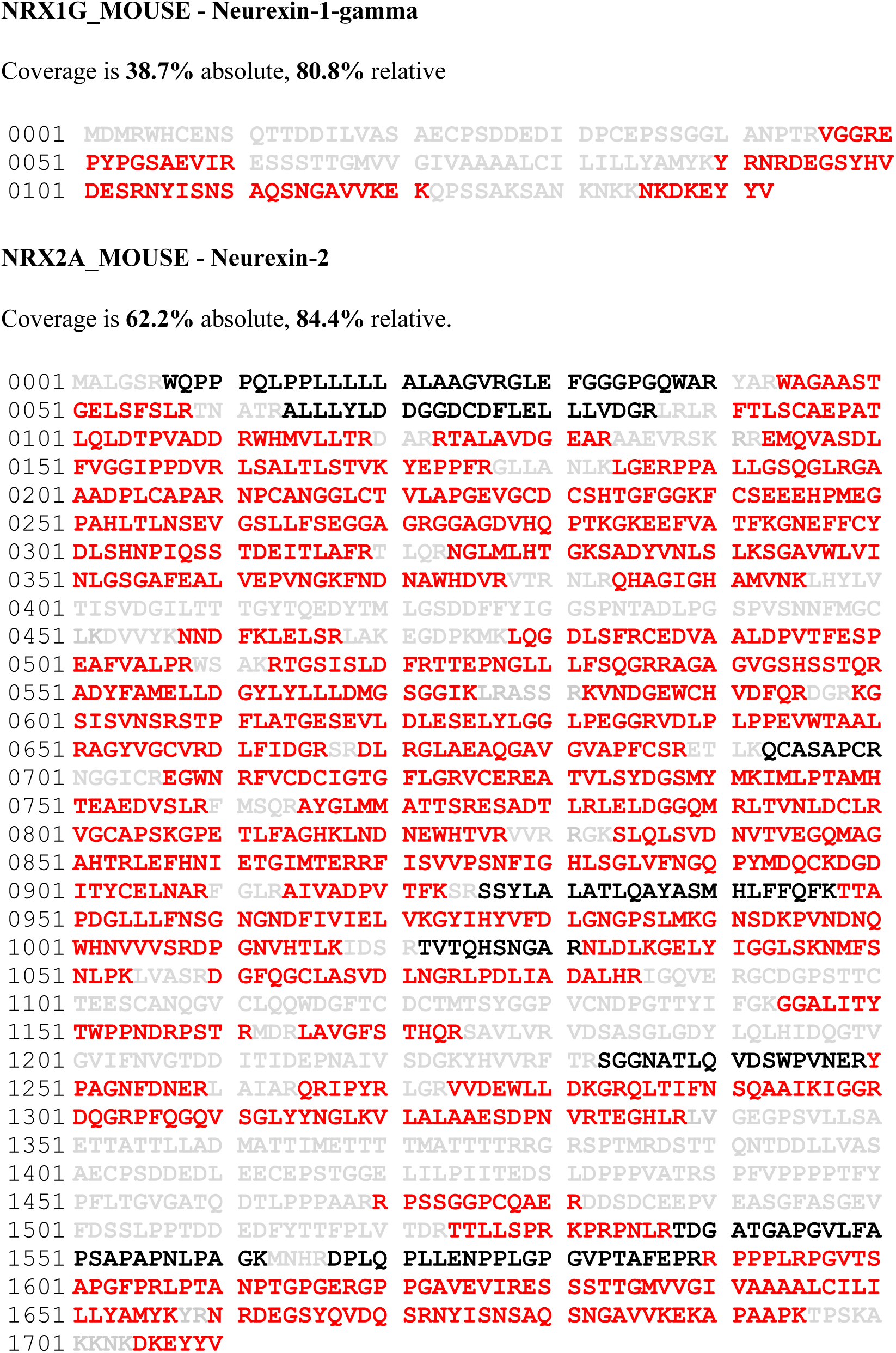
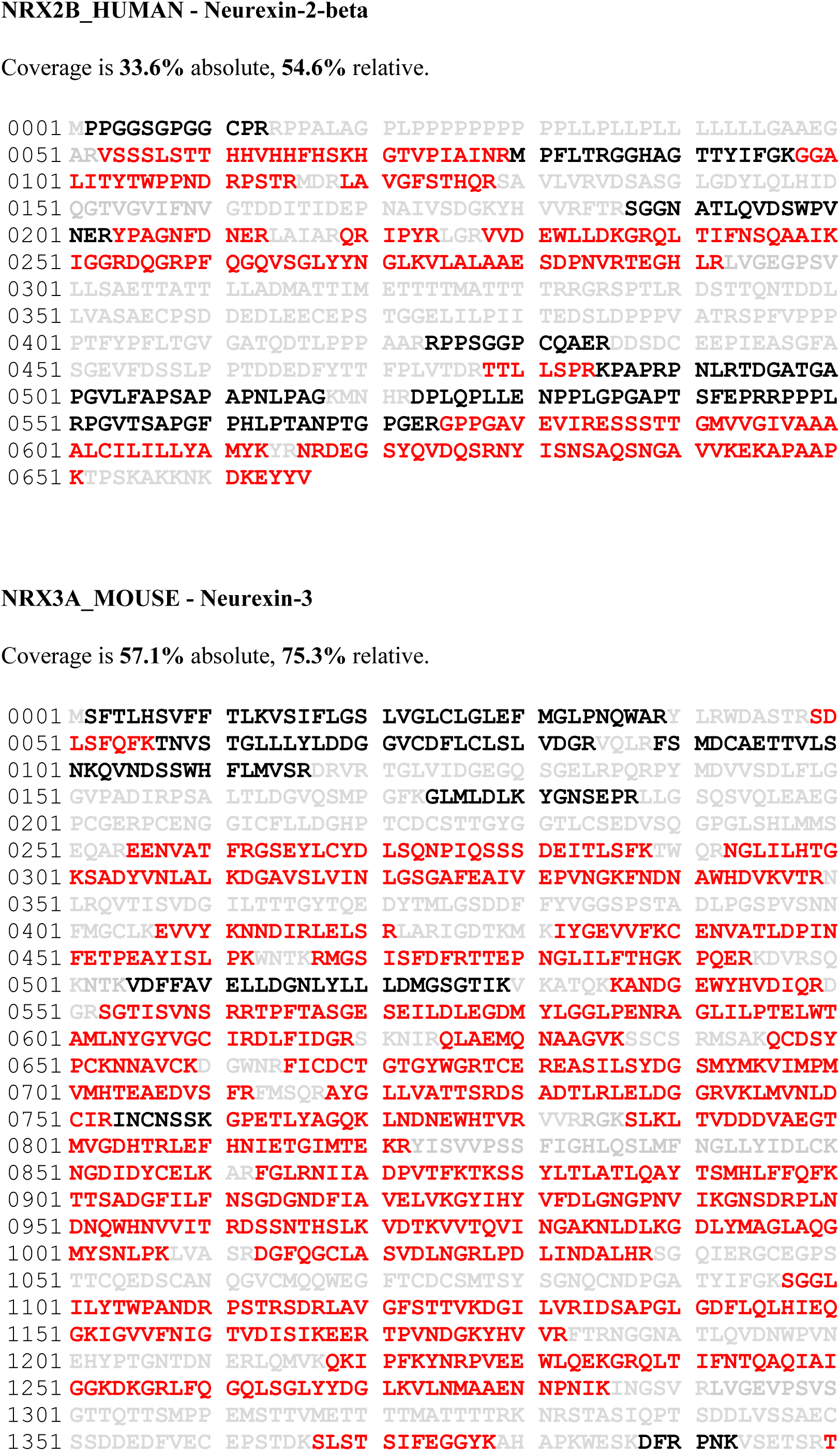
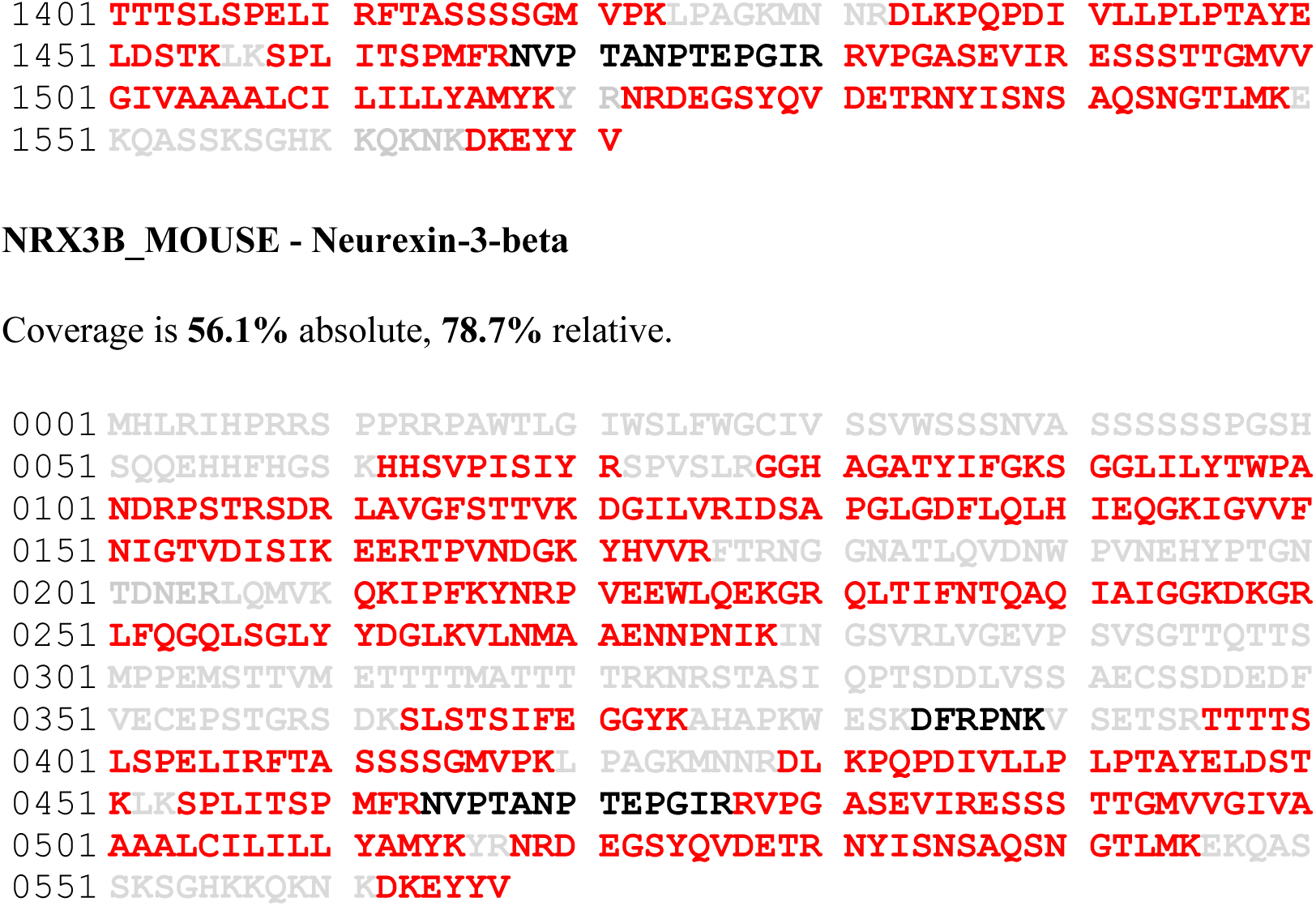
Coverage of primary sequences of NRX isoforms detected in pan-NRX1-3 APs. Peptides identified by mass spectrometry are in red; those accessible to, but not identified in MS/MS analyses are in black, and peptides hardly or not accessible to our MS/MS analyses (peptide mass >3 kDa or < 738 Da) are given in grey. Absolute coverage is ‘identified sequences / protein sequence’, relative coverage is ‘identified sequences / MS-accessible protein sequence’.

**Supplementary Figure 3.**
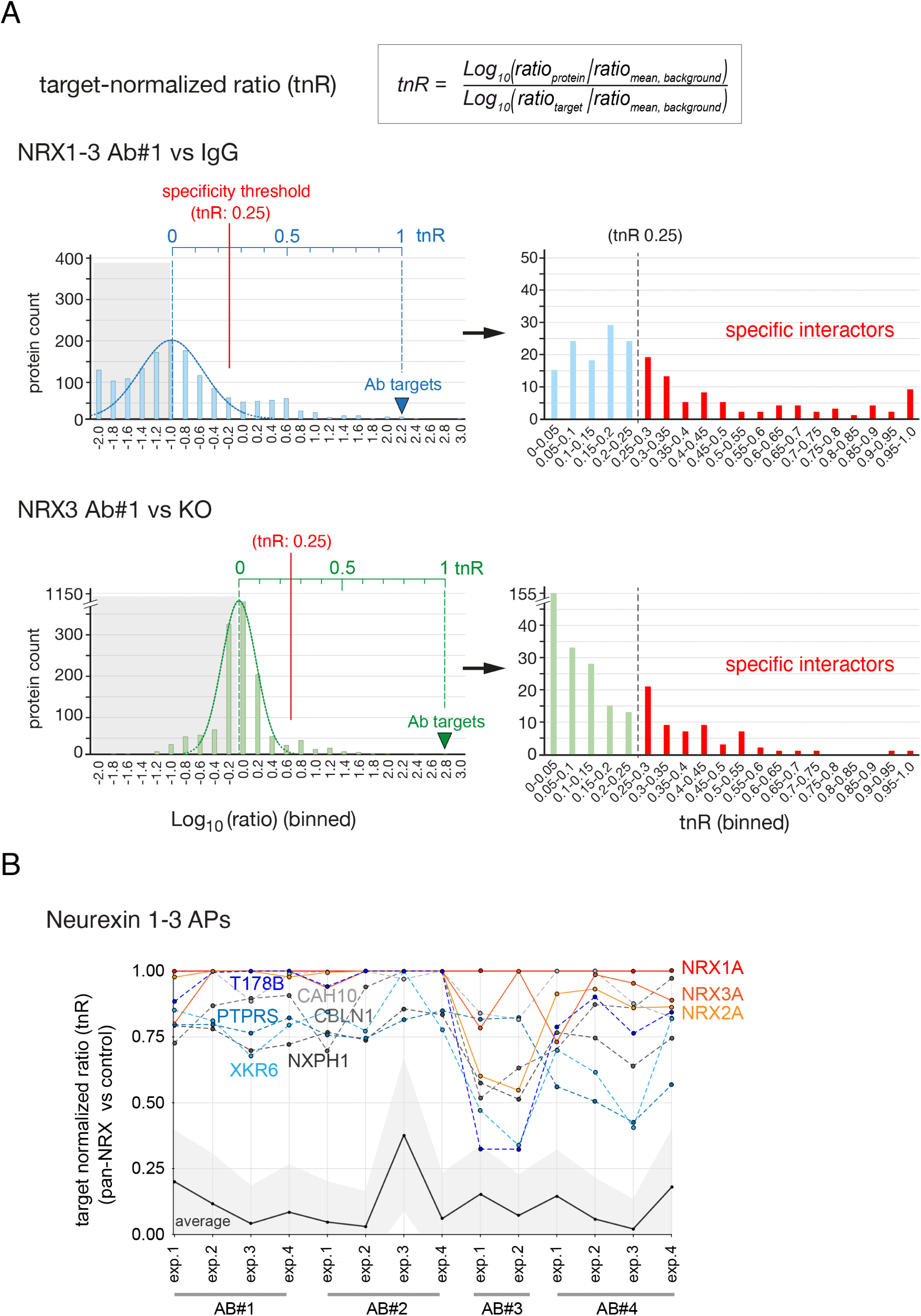
Evaluation of NRX APs by target-normalized ratios (tnR) (A), Top, equation for calculating target-normalized ratio values (tnRs, (Kocylowski et al., 2022)). Middle, example histograms depicting distribution of enrichment factors (i.e. abundance ratios of proteins in an anti-NRX1-3 AP versus an IgG control, log10-transformed) before (left) and after (right) re-scaling to tnRs with min = 0 derived from the gaussian distribution (dashed curve) of background proteins and max = 1 defined by the enrichment of the primary AP targets (NRX1A and NRX3A). Binning of ratios / tnR values as indicated (proteins with tnR = 0 were left out in the right plot for clarity). An empirical specificity threshold of tnR = 0.25 was applied resulting in 83 candidate interaction partners (highlighted red). Low, histograms and tnR-transformation as above for an anti-NRX3A AP from WT versus NRX3A-KO as specificity control yielding 63 candidate interactors. Note that, although target knockouts are generally superior antibody AP specificity controls, for the NRX antibodies used here numbers and distributions of background and co-enriched proteins are qualitatively similar. (B), Target-normalized ratio (tnR)-values of selected proteins specifically (co-)purified in 12 anti-NRX1-3 APs with four different anti-NRX1-3 ABs (see also Figure 1C). tnR-values >0.25 indicate specific co-purification (with the target). Average indicates average-tnRs calculated from all MS-identified proteins in the respective AP-eluate, grey area represents SDs of these average-tnRs.

**Supplementary Figure 4.**
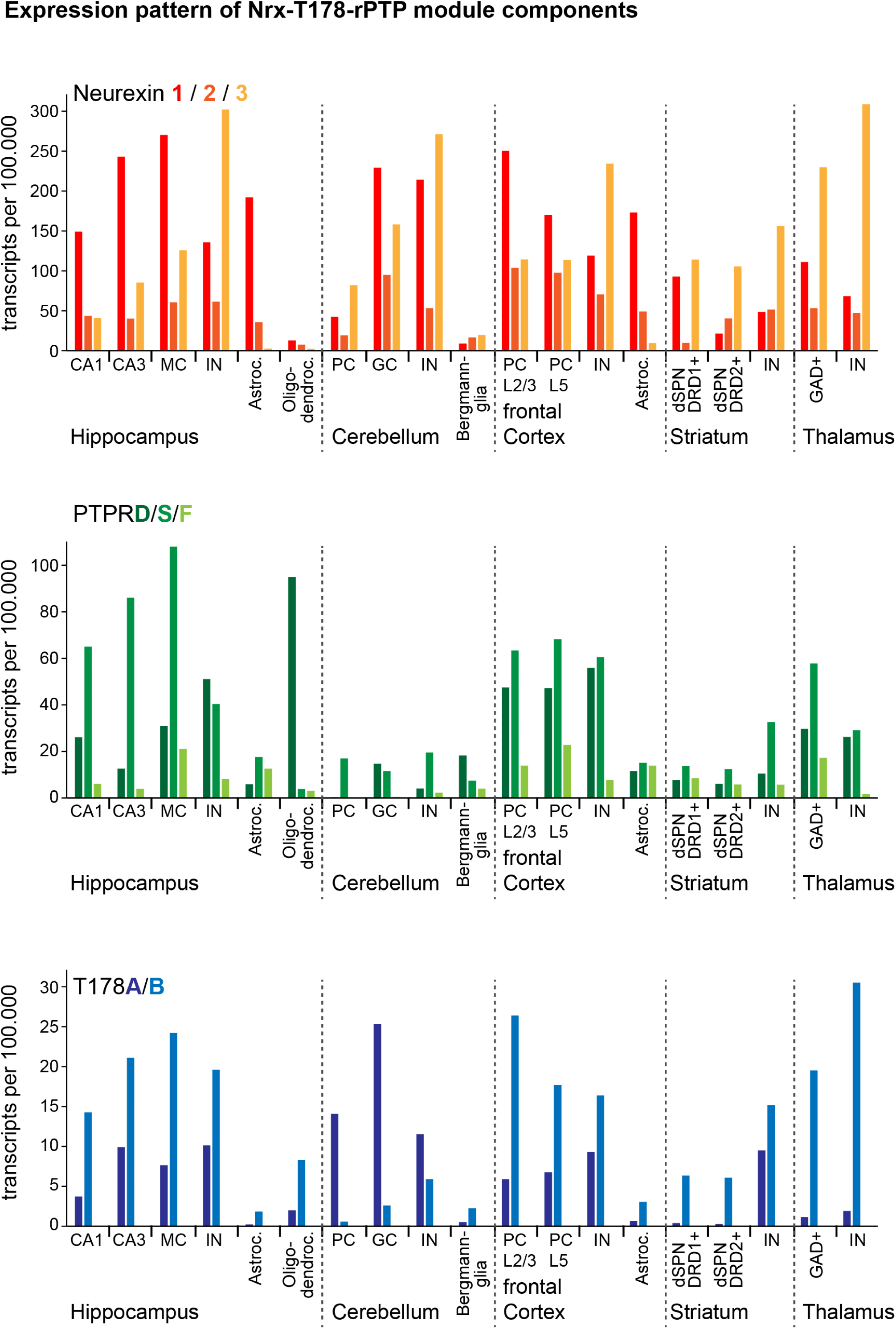
Expression pattern of NRX1-3, T178A/B and PTPRD/S/F in mouse brain. Transcriptome data were extracted from the DropViz data base (Saunders et al., 2018). Number of RNA transcripts per 100,000 unique molecular identifiers in the indicated brain regions and meta-groups according to DropViz single-cell transcriptomics data. CA1: CA1 pyramidal cells, CA3: CA3 pyramidal cells, DG: dentate gyrus, MC: mossy cells, IN: interneurons, GC: granule cells, PC: pyramidal cells, DRD1/2: dopamine receptor D1/2, GAD: glutamic acid decarboxylase. Note ubiquitous expression all of core-module constituents in both neurons and glia and distinct patterns for T178A (predominantly found in cerebellum).

**Supplementary Figure 5.**
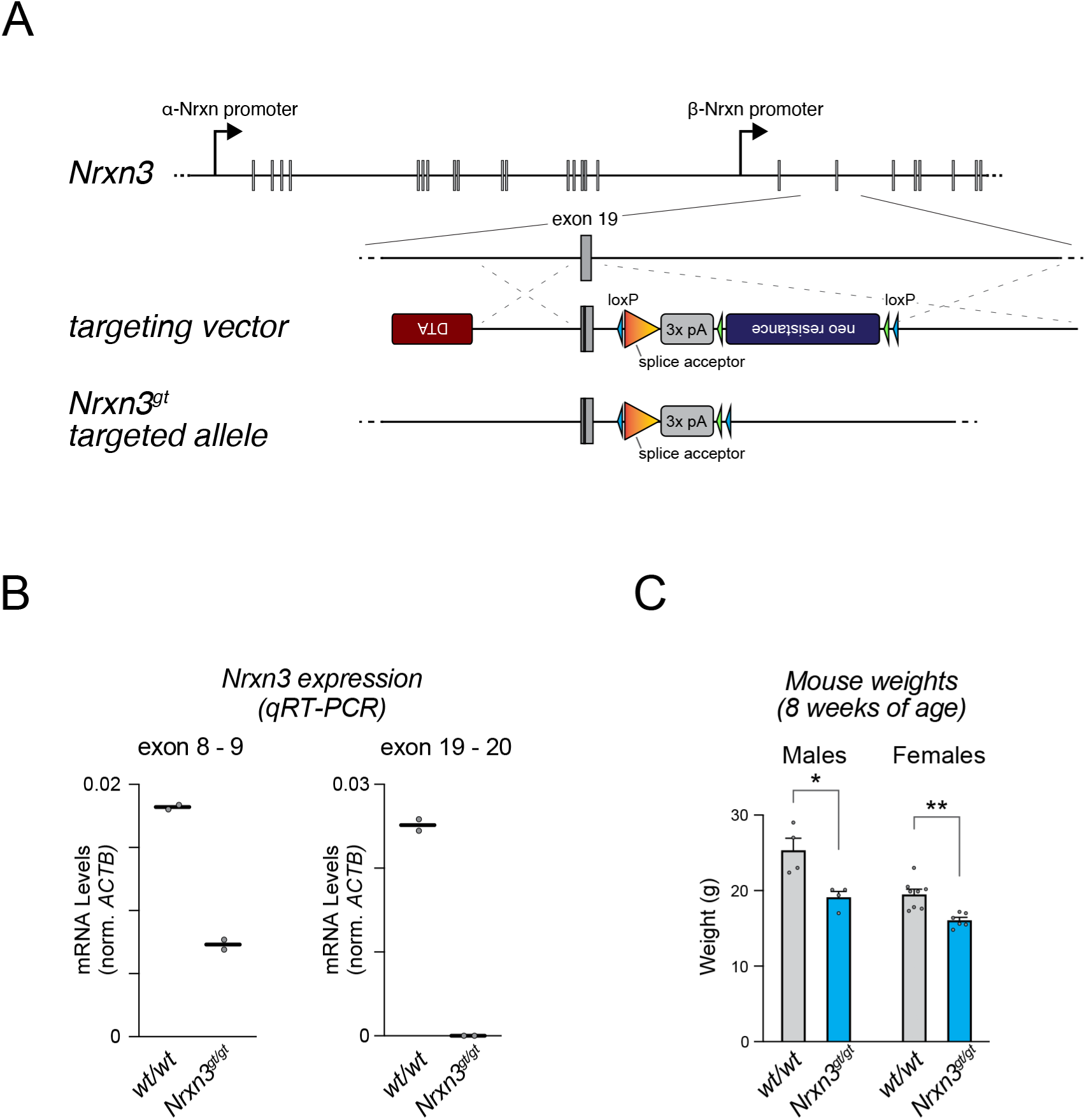
Design and validation of NRX3 KO mice. (A), Design of the targeting vector for gene-trap mutagenesis of NRX3 in mice. The gene-trap construct was introduced after the first exon common to both α- and β-isoforms. (B), Gene-trap insertion result in loss of mRNA. Levels of Nrxn3 transcripts before and after the targeted intron, respectively, monitored by qRT-PCR. **c**, Mouse weight at 8 weeks of age. *, p < 0.05 and **, p < 0.01 by two-tailed students t-test.

**Supplementary Figure 6.**
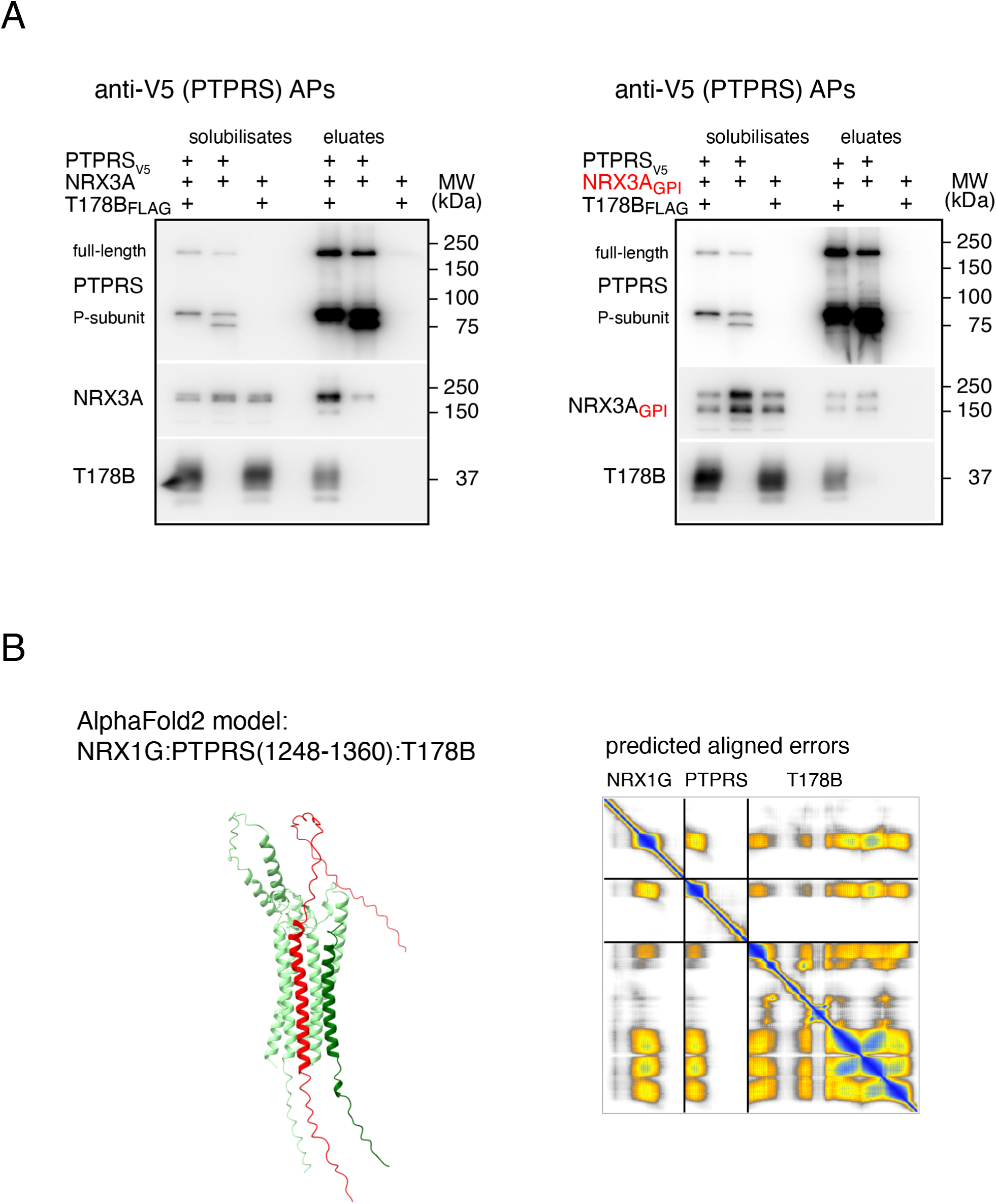
(A), **Reconstitution of NRX3A, T178B and PTPRS complexes** Western-probed SDS-PAGE separations of input and eluates of APs with anti-V5 ABs from tsA201 cells transiently expressing V5-tagged PTPRs, FLAG-tagged T178B and either NRX3A or a splice variant of NRX3A lacking the transmembrane domain and attached to the membrane via a GPI-anchor. Note that the presence of T178B appears critical for stable complex formation with NRX3A (similar as NRX1A, Figure 2E). In contrast, binding of NRX3A-GPI to PTPRS is weak and independent of the presence of T178B. (B), **Details of structural predictions by AlphaFold** Prediction of 3D-structural arrangement of the membrane-close region of the core-module composed of NRX1γ (mouse, aa 1-139, full length), PTPRS (mouse, aa 1251-1360) and T178B (human, 1-294, full length) by AlphaFold2-Multimer. Predicted aligned errors (PAE) provide estimates for the confidence of the correct position of two residues in the structure. Note high-confidence levels (colored blue) were only observed in the transmembrane regions; lower confidence levels are shown in yellow.

**Supplementary Figure 7.**
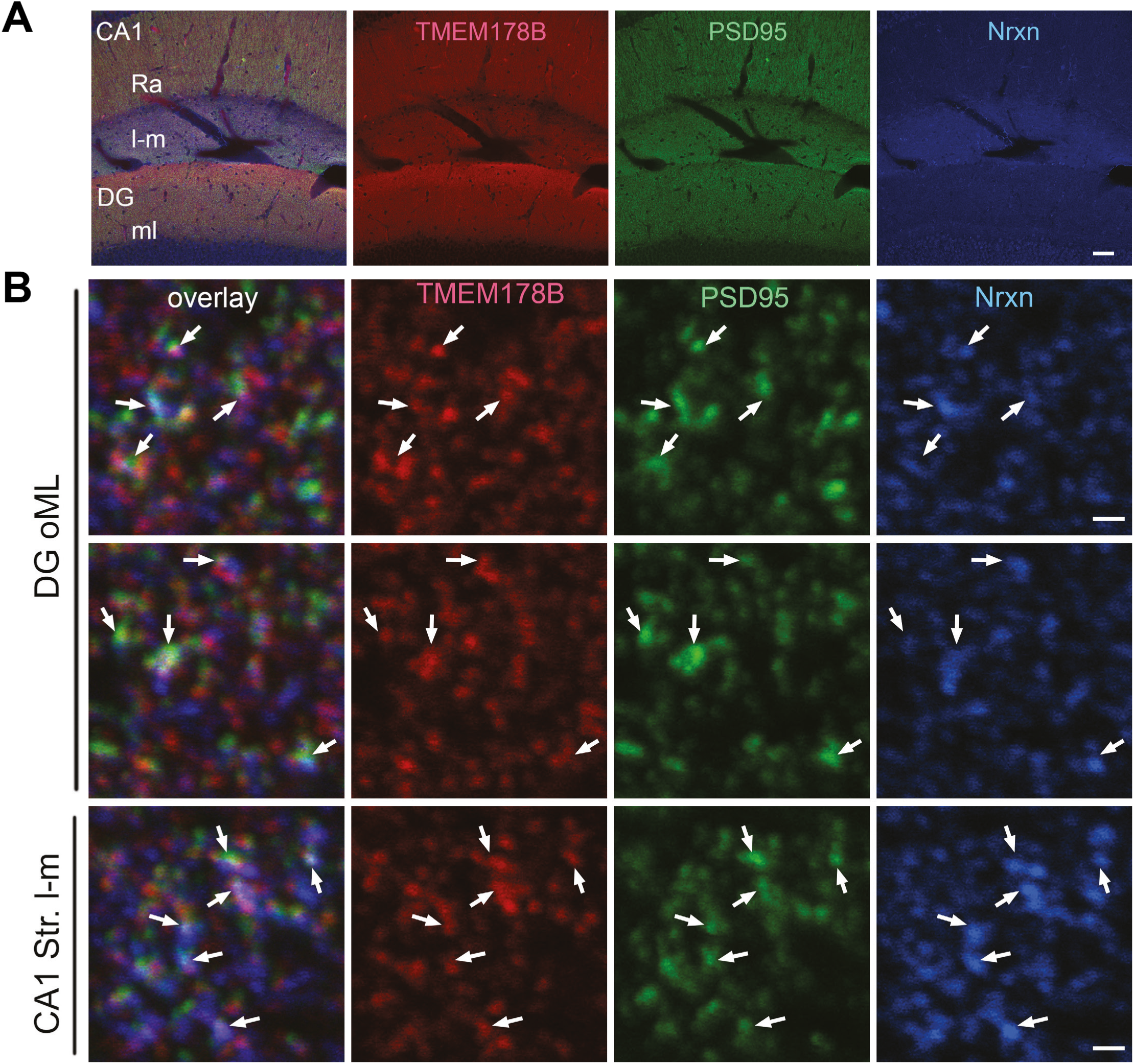
Localization of T178B to synapses together with NRX1-3 and PSD95/DLG4 in the hippocampus of adult mice. (A), A low-magnification view of the hippocampus of heterozygous T178B-HA mice shows that T178B (red), PSD95, and NRX are expressed in the stratum radiatum (Ra) and stratum lacunosum-moleculare (l-m) of the CA1 region, as well as in the molecular layer (ml) of the dentate gyrus (DG). Scale bar, 50 μm. (B), High-magnification views of the outer molecular layer of the dentate gyrus (DG oML) and the stratum lacunosum-moleculare (l-m) of CA1. Arrows indicate colocalization of T178B, PSD95, and NRX. Scale bar, 1 μm.

**Supplementary Figure 8.**
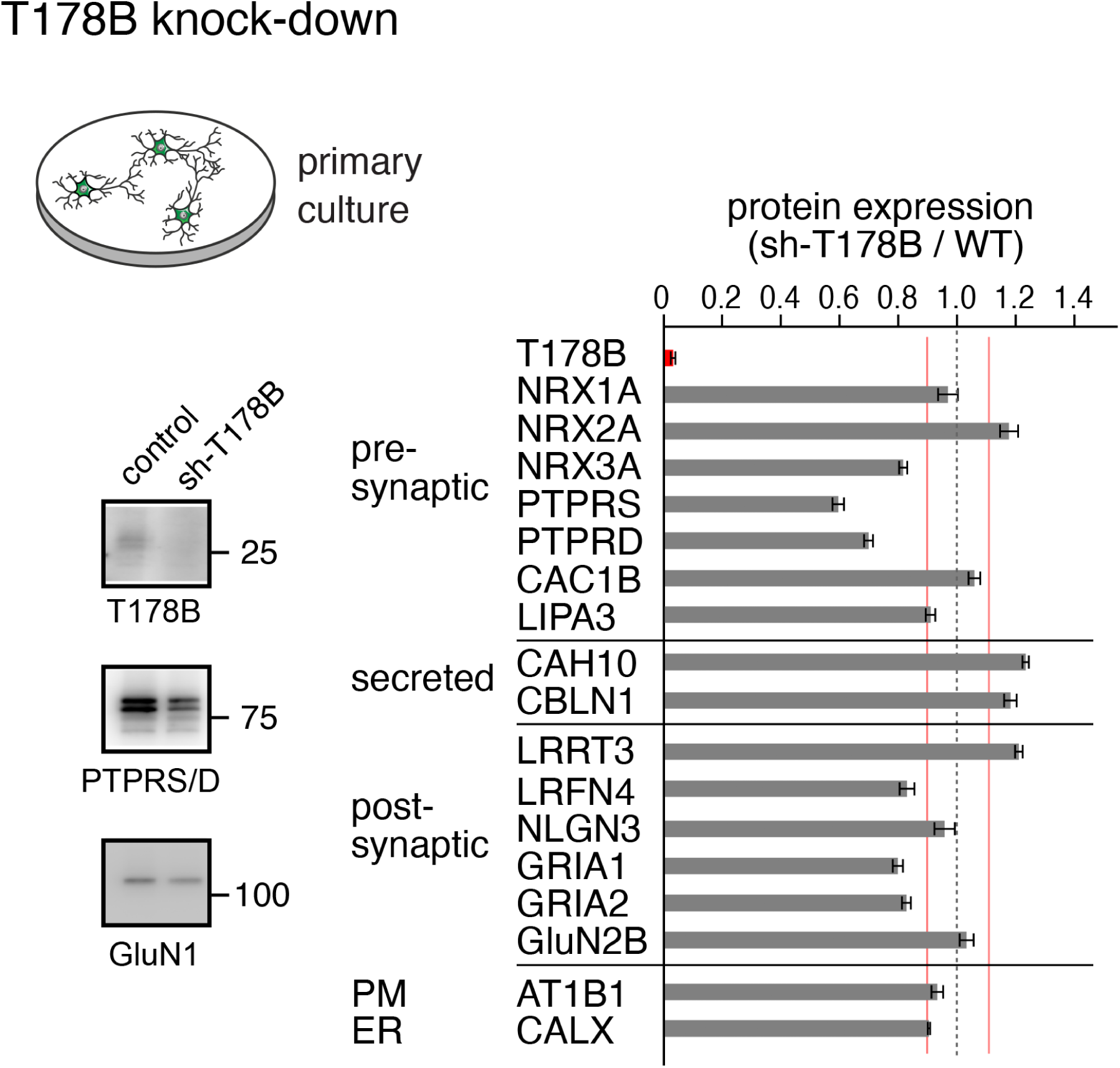
Ratio of protein abundances determined by quantitative MS in membranes from cultured cortical neurons infected either with AAV driving sh-T178B expression or with control virus. Note profound alteration in the expression of distinct synaptic proteins by sh-RNA-mediated knock-down of T178B: Expression of PTPRD, S was decreased by ∼40%, while essential subunits of NMDA receptors (GluN2B), Na/K-ATPase (AT1B1) and Calnexin (CALX) remained unchanged. Values are mean ± SEM of three samples of cultured neurons. Inset: Western-probed gel-separations indicating the effect of AAV:sh-T178B infection.

**Supplementary Figure 9.**
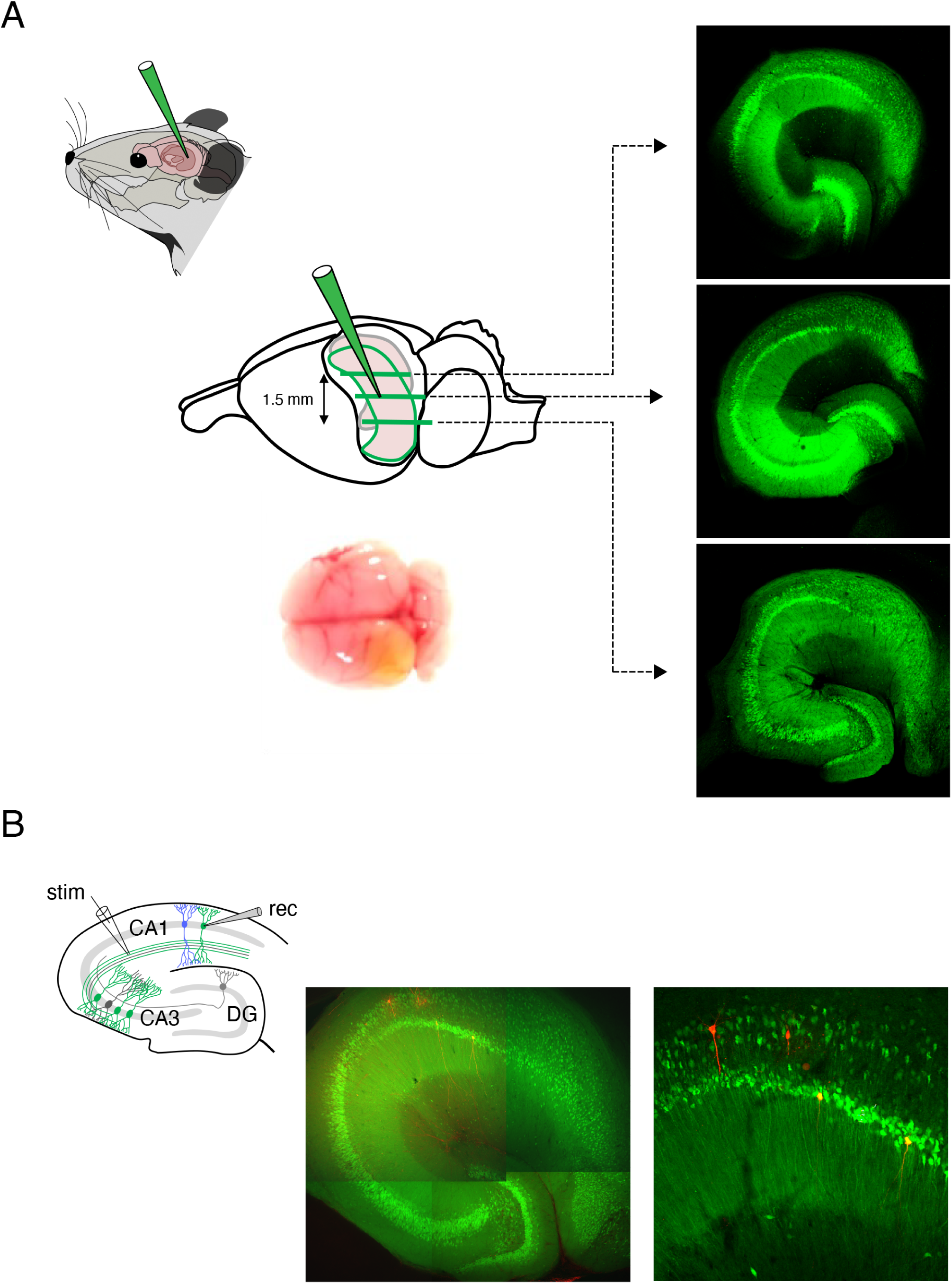
Stereotactic injection of AAV expressing sh-T178B into hippocampal CA3 regions and somatic recordings in CA1 pyramidal cells of adult mice. (A), Scheme depicting stereotactic injection into the hippocampal CA3 region and confocal images showing GFP-fluorescence in three brain slices to document spreading and transfection of cells by the AAV particles; GFP-expression is driven by the ubiquitin promotor (see generation of AAV-virus in Methods). Inset: Image showing unilateral injection into the left hemisphere. (B), Scheme illustrating slice-recordings of CA1 PCs in whole cell configuration and confocal images indicating four CA1 PCs filled with biocytin (red fluorescence) through the patch-pipette during recordings. Note two of these cells were infected by the AAV (yellow fluorescence), two cells were non-infected (red fluorescence).

**Supplementary Figure 10.**
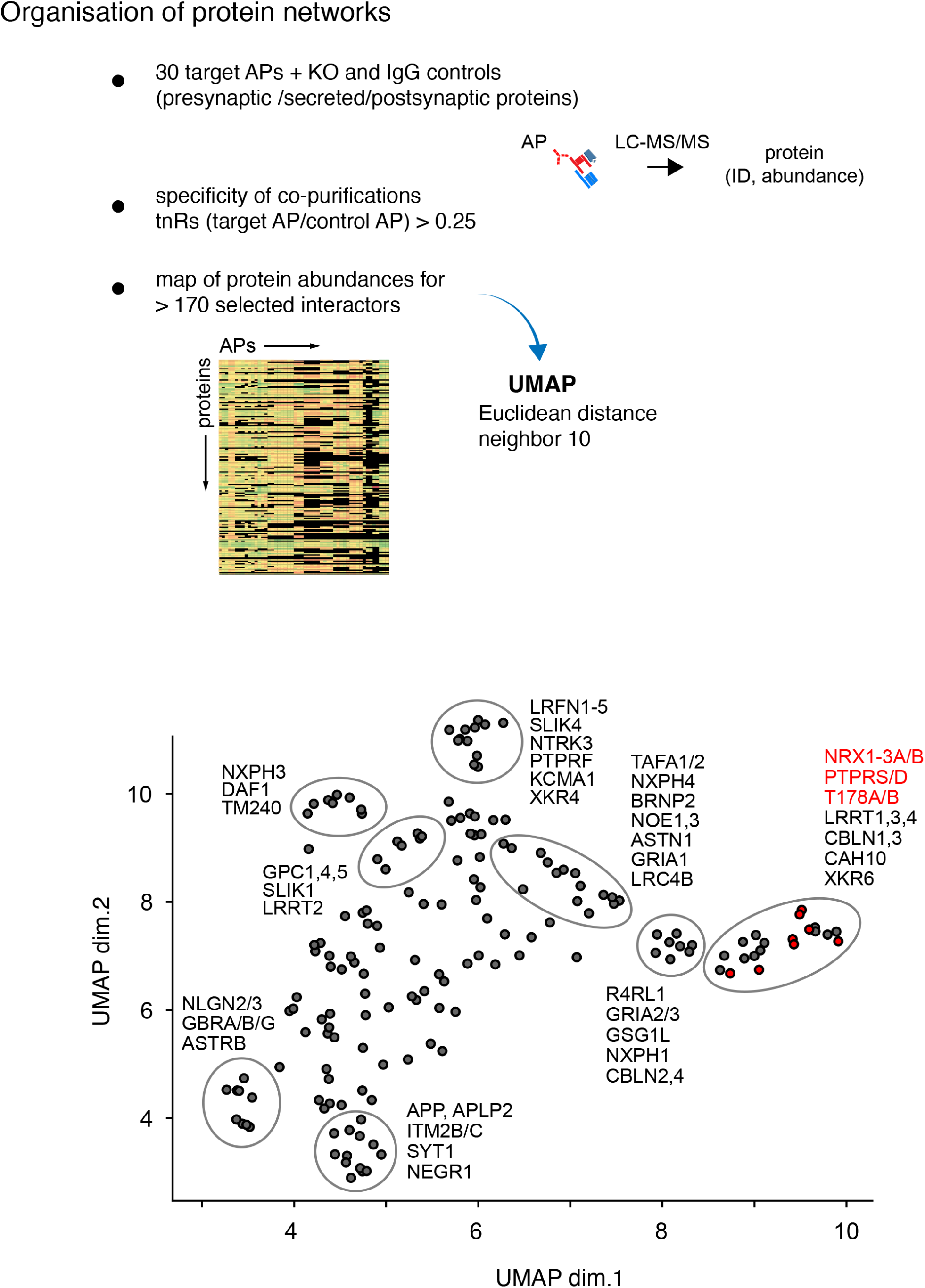
UMAP (Uniform Manifold Approximation and Projection, (Sainburg et al., 2021)) cluster-analysis performed with the MS-derived abundance values of all proteins (total of 172) that were specifically co-purified in NRX1-3 and reverse APs with a total of 30 different target proteins. UMAP-derived subclusters (of similar behavior over the entire set of APs) were encircled with selected constituents annotated.

### Supplementary Tables

**Supplementary Table 1**

Characterization of the anti-NRX ABs used by MS-analysis of respective target APs (related to Figures 1C-E, Table S2)

**Supplementary Table 2**

tnR-values of APs with the anti-NRX1-3 ABs (**Table 1**), and with the sub-type specific anti-NRX1, 2, 3 ABs

**Supplementary Table 3**

tnR-values of APs with anti-NRX1-3 and anti-PTPRs ABs from ER-membranes

**Supplementary Table 4**

Protein abundance ratios determined in membrane fractions from cortical cultures infected with AAV: sh-T178B or control AAV: GFP

**Supplementary Table 5**

tnR-values of specifically co-purified proteins in the target APs indicated in Figure 5

## References

1. Südhof TC. Cell. 2017;171(4):745–69.

2. Sterky FH et al. Proc Natl Acad Sci U S A. 2017;114(7):E1253–62.

3. Aydin D et al. Exp brain Res. 2012;217(3–4):423–34.

4. Wang J, et al. Cell. 2021;184(24):5869–5885.e25.

5. Fishell G et al. Development 1991; 113(3):755–65.

6. Sandhu J, et al. Cell 2018 ;175(2):514–529.e20.

7. Terashima M et al. J Neurosci Res 2010;88(7):1387–93.

8. Matsuda K et al. Neuron 2016;90(4):752–67.

9. Frei JA, Stoeckli ET. Mol Cell Neurosci. 2017;81:41–8.

10. Michalak M. J Cell Mol Med. 2024;28(5).

11. Uemura T et al. Cell 2010;141(6):1068–79.

12. Südhof TC. Curr Opin Neurobiol. 2023 Aug 1;81

13. Silkensen JR et al. J Clin Invest 1995;96(6):2646.

14. Dityatev A et al. Neuron Glia Biol 2008;4(3):197–209.

15. Cull-Candy S et al. Curr Opin Neurobiol 2006;16(3):288–97.

16. Ramos C et al. J Biol Chem. 2012;287(24):20176–86.

17. Steegmaler M et al. Nature;373(6515):615–20.

18. Matsuda S et al. J Biol Chem. 2005;280(32):28912–6.

19. Dinamarca MC et al. Nat Commun. 2019;10(1).

20. Nwaobi SE et al. Acta Neuropathol 2016;132(1).

21. Griguoli M et al. J Physiol. 2016;594(13):3489–500.

22. Woo J et al. Nat Neurosci. 2009 Apr;12(4):428–37.

23. Lie E et al. Front Mol Neurosci 2018;11:105.

24. Soler-Llavina GJ et al. Neuron 2013;79(3):439–46.

25. Um JW et al. Cell Rep 2016;14(4):808–22.

26. DeWit J et al. Neuron 2013;79(4):696–711.

27. Laboute T et al. Science 2023;379(6639):1352–8.

28. Varoqueaux F et al. Neuron 2006;51(6):741–54.

29. Debaigt C et al. J Biol Chem 2004;279(34):35687–91.

30. Missler M et al. J Biol Chem 1998;273(52):34716–23.

31. Lu WC et al. Neural Regen Res 2018;13(3):427–33.

32. Matt L et al. Cell Rep. 2018;22(9):2246–53.

33. Cornejo F et al. Front cell Dev Biol 2021;9.

34. Yoo SW et al. J Comp Neurol 2017;525(2):291–301.

35. Nguyen-Ba-Charvet KT, Chédotal A. J Physiol Paris 2002;96(1–2):91–8.

36. Pan Q, et al. Gastroenterology 2018;155(5):1578–1592.e16.

37. Trovo L, et al EMBO Rep 2024 (in press)

38. Li B et al. Development 2022;149(17).

39. Khalaj AJ et al. J Cell Biol. 2020;219(9).

40. Kojima I, Nagasawa M. Handb Exp Pharmacol 2014;222:247–72.

41. Suzuki J et al. J Biol Chem 2014;289(44):30257–67.

## References

Anderson, G.R., Aoto, J., Tabuchi, K., Foldy, C., Covy, J., Yee, A.X., Wu, D., Lee, S.J., Chen, L., Malenka, R.C., and Sudhof, T.C. (2015). beta-Neurexins Control Neural Circuits by Regulating Synaptic Endocannabinoid Signaling. Cell 162, 593–606. 10.1016/j.cell.2015.06.056.

Althof, D., Baehrens, D., Watanabe, M., Suzuki, N., Fakler, B., and Kulik, A. (2015). Inhibitory and excitatory axon terminals share a common nano-architecture of their Cav2.1 (P/Q-type) Ca^2+^ channels. Front Cell Neurosci 9, 315. 10.3389/fncel.2015.00315.

Aoto, J., Foldy, C., Ilcus, S.M., Tabuchi, K., and Sudhof, T.C. (2015). Distinct circuit-dependent functions of presynaptic neurexin-3 at GABAergic and glutamatergic synapses. Nat Neurosci 18, 997–1007. 10.1038/nn.4037.

Biederer, T., Kaeser, P.S., and Blanpied, T.A. (2017). Transcellular Nanoalignment of Synaptic Function. Neuron 96, 680–696. 10.1016/j.neuron.2017.10.006.

Bildl, W., Haupt, A., Muller, C.S., Biniossek, M.L., Thumfart, J.O., Huber, B., Fakler, B., and Schulte, U. (2012). Extending the dynamic range of label-free mass spectrometric quantification of affinity purifications. Mol Cell Proteomics 11, M111 007955. 10.1074/mcp.M111.007955.

Boudkkazi, S., Brechet, A., Schwenk, J., and Fakler, B. (2014). Cornichon2 dictates the time course of excitatory transmission at individual hippocampal synapses. Neuron 82, 848–858. 10.1016/j.neuron.2014.03.031.

Boudkkazi, S., Schwenk, J., Nakaya, N., Brechet, A., Kollewe, A., Harada, H., Bildl, W., Kulik, A., Dong, L., Sultana, A., et al. (2023). A Noelin-organized extracellular network of proteins required for constitutive and context-dependent anchoring of AMPA-receptors. Neuron 111, 2544–2556 e2549. 10.1016/j.neuron.2023.07.013.

Chen, L.Y., Jiang, M., Zhang, B., Gokce, O., and Sudhof, T.C. (2017). Conditional Deletion of All Neurexins Defines Diversity of Essential Synaptic Organizer Functions for Neurexins. Neuron 94, 611–625 e614. 10.1016/j.neuron.2017.04.011.

Cox, J., Hein, M.Y., Luber, C.A., Paron, I., Nagaraj, N., and Mann, M. (2014). Accurate proteome-wide label-free quantification by delayed normalization and maximal peptide ratio extraction, termed MaxLFQ. Mol Cell Proteomics 13, 2513–2526. 10.1074/mcp.M113.031591.

Cvetkovska, V., Ge, Y., Xu, Q., Li, S., Zhang, P., and Craig, A.M. (2022). Neurexin-beta Mediates the Synaptogenic Activity of Amyloid Precursor Protein. J Neurosci 42, 8936–8947. 10.1523/JNEUROSCI.0511-21.2022.

Dai, Y., Taru, H., Deken, S.L., Grill, B., Ackley, B., Nonet, M.L., and Jin, Y. (2006). SYD-2 Liprin-alpha organizes presynaptic active zone formation through ELKS. Nat Neurosci 9, 1479–1487. 10.1038/nn1808.

de Wit, J., and Ghosh, A. (2016). Specification of synaptic connectivity by cell surface interactions. Nat Rev Neurosci 17, 22–35. 10.1038/nrn.2015.3.

de Wit, J., Sylwestrak, E., O’Sullivan, M.L., Otto, S., Tiglio, K., Savas, J.N., Yates, J.R., 3rd, Comoletti, D., Taylor, P., and Ghosh, A. (2009). LRRTM2 interacts with Neurexin1 and regulates excitatory synapse formation. Neuron 64, 799–806. 10.1016/j.neuron.2009.12.019.

de Wit, J., O’Sullivan, M.L., Savas, J.N., Condomitti, G., Caccese, M.C., Vennekens, K.M., Yates, J.R., and Ghosh, A. (2013). Unbiased discovery of glypican as a receptor for LRRTM4 in regulating excitatory synapse development. Neuron 79, 696–711. 10.1016/J.NEURON.2013.06.049.

Emperador-Melero, J., Andersen, J.W., Metzbower, S.R., Levy, A.D., Dharmasri, P.A., de Nola, G., Blanpied, T.A., and Kaeser, P.S. (2024). Distinct active zone protein machineries mediate Ca^2+^ channel clustering and vesicle priming at hippocampal synapses. Nat Neurosci 27, 1680–1694. 10.1038/s41593-024-01720-5.

Gomez, A.M., Traunmuller, L., and Scheiffele, P. (2021). Neurexins: molecular codes for shaping neuronal synapses. Nat Rev Neurosci 22, 137–151. 10.1038/s41583-020-00415-7.

Graf, E.R., Zhang, X., Jin, S.X., Linhoff, M.W., and Craig, A.M. (2004). Neurexins induce differentiation of GABA and glutamate postsynaptic specializations via neuroligins. Cell 119, 1013–1026. 10.1016/j.cell.2004.11.035.

Hauser, D., Behr, K., Konno, K., Schreiner, D., Schmidt, A., Watanabe, M., Bischofberger, J., and Scheiffele, P. (2022). Targeted proteoform mapping uncovers specific Neurexin-3 variants required for dendritic inhibition. Neuron 110, 2094–2109 e2010. 10.1016/j.neuron.2022.04.017.

Horn, K.E., Xu, B., Gobert, D., Hamam, B.N., Thompson, K.M., Wu, C.L., Bouchard, J.F., Uetani, N., Racine, R.J., Tremblay, M.L., et al. (2012). Receptor protein tyrosine phosphatase sigma regulates synapse structure, function and plasticity. J Neurochem 122, 147–161. 10.1111/j.1471-4159.2012.07762.x.

Itzhak, D.N., Tyanova, S., Cox, J., and Borner, G.H. (2016). Global, quantitative and dynamic mapping of protein subcellular localization. Elife 5. 10.7554/eLife.16950.

Jang, S., Lee, H., and Kim, E. (2017). Synaptic adhesion molecules and excitatory synaptic transmission. Curr Opin Neurobiol 45, 45–50. 10.1016/j.conb.2017.03.005.

Khalaj, A.J., Sterky, F.H., Sclip, A., Schwenk, J., Brunger, A.T., Fakler, B., and Sudhof, T.C. (2020). Deorphanizing FAM19A proteins as pan-neurexin ligands with an unusual biosynthetic binding mechanism. J Cell Biol 219, e202004164. 10.1083/jcb.202004164.

Kocylowski, M.K., Aypek, H., Bildl, W., Helmstadter, M., Trachte, P., Dumoulin, B., Wittosch, S., Kuhne, L., Aukschun, U., Teetzen, C., et al. (2022). A slit-diaphragm-associated protein network for dynamic control of renal filtration. Nat Commun 13, 6446. 10.1038/s41467-022-33748-1.

Kollewe, A., Chubanov, V., Tseung, F.T., Correia, L., Schmidt, E., Rossig, A., Zierler, S., Haupt, A., Muller, C.S., Bildl, W., et al. (2021). The molecular appearance of native TRPM7 channel complexes identified by high-resolution proteomics. Elife 10. 10.7554/eLife.68544.

Kollewe, A., Schwarz, Y., Oleinikov, K., Raza, A., Haupt, A., Wartenberg, P., Wyatt, A., Boehm, U., Ectors, F., Bildl, W., et al. (2022). Subunit composition, molecular environment, and activation of native TRPC channels encoded by their interactomes. Neuron 110, 4162–4175 e4167. 10.1016/j.neuron.2022.09.029.

Kusch, V., Bornschein, G., Loreth, D., Bank, J., Jordan, J., Baur, D., Watanabe, M., Kulik, A., Heckmann, M., Eilers, J., and Schmidt, H. (2018). Munc13-3 Is Required for the Developmental Localization of Ca^2+^ Channels to Active Zones and the Nanopositioning of Cav2.1 Near Release Sensors. Cell Rep 22, 1965–1973. 10.1016/j.celrep.2018.02.010.

Krueger-Burg, D., Papadopoulos, T., and Brose, N. (2017). Organizers of inhibitory synapses come of age. Curr Opin Neurobiol 45, 66–77. 10.1016/j.conb.2017.04.003.

Laboute, T., Zucca, S., Holcomb, M., Patil, D.N., Garza, C., Wheatley, B.A., Roy, R.N., Forli, S., and Martemyanov, K.A. (2023). Orphan receptor GPR158 serves as a metabotropic glycine receptor: mGlyR. Science 379, 1352–1358. 10.1126/science.add7150.

Li, B., Brusman, L., Dahlka, J., and Niswander, L.A. (2022). TMEM132A ensures mouse caudal neural tube closure and regulates integrin-based mesodermal migration. Development 149. 10.1242/dev.200442.

Lie, E., Li, Y., Kim, R., and Kim, E. (2018). SALM/Lrfn Family Synaptic Adhesion Molecules. Frontiers in molecular neuroscience 11, 105–105. 10.3389/fnmol.2018.00105.

Marco de la Cruz, B., Campos, J., Molinaro, A., Xie, X., Jin, G., Wei, Z., Acuna, C., and Sterky, F.H. (2024). Liprin-alpha proteins are master regulators of human presynapse assembly. Nat Neurosci 27, 629–642. 10.1038/s41593-024-01592-9.

Martin-Belmonte, A., Aguado, C., Alfaro-Ruiz, R., Kulik, A., de la Ossa, L., Moreno-Martinez, A.E., Alberquilla, S., Garcia-Carracedo, L., Fernandez, M., Fajardo-Serrano, A., et al. (2025). Nanoarchitecture of Ca(V)2.1 channels and GABA(B) receptors in the mouse hippocampus: Impact of APP/PS1 pathology. Brain Pathol 35, e13279. 10.1111/bpa.13279.

Matsuda, K., and Yuzaki, M. (2011). Cbln family proteins promote synapse formation by regulating distinct neurexin signaling pathways in various brain regions. Eur J Neurosci 33, 1447–1461. 10.1111/j.1460-9568.2011.07638.x.

McInnes, L., Healy, J., Saul, N., and Großberger, L. (2018). UMAP: Uniform Manifold Approximation and Projection. J Open Source Software 3, 861.

Missler, M., and Sudhof, T.C. (1998). Neurexophilins form a conserved family of neuropeptide-like glycoproteins. J Neurosci 18, 3630–3638. 10.1523/JNEUROSCI.18-10-03630.1998.

Montoliu-Gaya, L., Tietze, D., Kaminski, D., Mirgorodskaya, E., Tietze, A.A., and Sterky, F.H. (2021). CA10 regulates neurexin heparan sulfate addition via a direct binding in the secretory pathway. EMBO Rep 22, e51349. 10.15252/embr.202051349.

Muhammad, K., Reddy-Alla, S., Driller, J.H., Schreiner, D., Rey, U., Bohme, M.A., Hollmann, C., Ramesh, N., Depner, H., Lutzkendorf, J., et al. (2015). Presynaptic spinophilin tunes neurexin signalling to control active zone architecture and function. Nat Commun 6, 8362. 10.1038/ncomms9362.

Muller, C.S., Haupt, A., Bildl, W., Schindler, J., Knaus, H.G., Meissner, M., Rammner, B., Striessnig, J., Flockerzi, V., Fakler, B., and Schulte, U. (2010). Quantitative proteomics of the Cav2 channel nano-environments in the mammalian brain. Proc Natl Acad Sci U S A 107, 14950–14957. 10.1073/pnas.1005940107.

Muller, C.S., Bildl, W., Klugbauer, N., Haupt, A., Fakler, B., and Schulte, U. (2019). High-Resolution Complexome Profiling by Cryoslicing BN-MS Analysis. J Vis Exp. 10.3791/60096.

Neniskyte, U., Kuliesiute, U., Vadisiute, A., Jevdokimenko, K., Coletta, L., Deivasigamani, S., Pamedytyte, D., Daugelaviciene, N., Dabkeviciene, D., Perlas, E., et al. (2023). Phospholipid scramblase Xkr8 is required for developmental axon pruning via phosphatidylserine exposure. EMBO J 42, e111790. 10.15252/embj.2022111790.

Nozawa, K., Hayashi, A., Motohashi, J., Takeo, Y.H., Matsuda, K., and Yuzaki, M. (2018). Cellular and Subcellular Localization of Endogenous Neuroligin-1 in the Cerebellum. Cerebellum 17, 709–721. 10.1007/s12311-018-0966-x.

Nozawa, K., Sogabe, T., Hayashi, A., Motohashi, J., Miura, E., Arai, I., and Yuzaki, M. (2022). In vivo nanoscopic landscape of neurexin ligands underlying anterograde synapse specification. Neuron 110, 3168–3185 e3168. 10.1016/j.neuron.2022.07.027.

Owald, D., Khorramshahi, O., Gupta, V.K., Banovic, D., Depner, H., Fouquet, W., Wichmann, C., Mertel, S., Eimer, S., Reynolds, E., et al. (2012). Cooperation of Syd-1 with Neurexin synchronizes pre-with postsynaptic assembly. Nat Neurosci 15, 1219–1226. 10.1038/nn.3183.

Roppongi, R.T., Dhume, S.H., Padmanabhan, N., Silwal, P., Zahra, N., Karimi, B., Bomkamp, C., Patil, C.S., Champagne-Jorgensen, K., Twilley, R.E., et al. (2020). LRRTMs Organize Synapses through Differential Engagement of Neurexin and PTPsigma. Neuron 106, 108–125 e112. 10.1016/j.neuron.2020.01.003.

Sainburg, T., McInnes, L., and Gentner, T.Q. (2021). Parametric UMAP Embeddings for Representation and Semisupervised Learning. Neural Comput 33, 2881–2907. 10.1162/neco_a_01434.

Saunders, A., Macosko, E.Z., Wysoker, A., Goldman, M., Krienen, F.M., de Rivera, H., Bien, E., Baum, M., Bortolin, L., Wang, S., et al. (2018). Molecular Diversity and Specializations among the Cells of the Adult Mouse Brain. Cell 174, 1015–1030 e1016. 10.1016/j.cell.2018.07.028.

Scheiffele, P., Fan, J., Choih, J., Fetter, R., and Serafini, T. (2000). Neuroligin expressed in nonneuronal cells triggers presynaptic development in contacting axons. Cell 101, 657–669. 10.1016/s0092-8674(00)80877-6.

Schmidt, N., Kollewe, A., Constantin, C.E., Henrich, S., Ritzau-Jost, A., Bildl, W., Saalbach, A., Hallermann, S., Kulik, A., Fakler, B., and Schulte, U. (2017). Neuroplastin and Basigin Are Essential Auxiliary Subunits of Plasma Membrane Ca^2+^-ATPases and Key Regulators of Ca^2+^ Clearance. Neuron 96, 827–838 e829. 10.1016/j.neuron.2017.09.038.

Schreiner, D., Nguyen, T.M., Russo, G., Heber, S., Patrignani, A., Ahrne, E., and Scheiffele, P. (2014). Targeted combinatorial alternative splicing generates brain region-specific repertoires of neurexins. Neuron 84, 386–398. 10.1016/j.neuron.2014.09.011.

Schulte, U., den Brave, F., Haupt, A., Gupta, A., Song, J., Muller, C.S., Engelke, J., Mishra, S., Martensson, C., Ellenrieder, L., et al. (2023). Mitochondrial complexome reveals quality-control pathways of protein import. Nature 614, 153–159. 10.1038/s41586-022-05641-w.

Schwenk, J., Boudkkazi, S., Kocylowski, M.K., Brechet, A., Zolles, G., Bus, T., Costa, K., Kollewe, A., Jordan, J., Bank, J., et al. (2019). An ER Assembly Line of AMPA-Receptors Controls Excitatory Neurotransmission and Its Plasticity. Neuron 104, 680–692 e689. 10.1016/j.neuron.2019.08.033.

Schwenk, J., and Fakler, B. (2021). Building of AMPA-type glutamate receptors in the endoplasmic reticulum and its implication for excitatory neurotransmission. J Physiol 599, 2639–2653. 10.1113/JP279025.

Schwenk, J., Harmel, N., Brechet, A., Zolles, G., Berkefeld, H., Muller, C.S., Bildl, W., Baehrens, D., Huber, B., Kulik, A., et al. (2012). High-resolution proteomics unravel architecture and molecular diversity of native AMPA receptor complexes. Neuron 74, 621–633. 10.1016/j.neuron.2012.03.034.

Schwenk, J., Metz, M., Zolles, G., Turecek, R., Fritzius, T., Bildl, W., Tarusawa, E., Kulik, A., Unger, A., Ivankova, K., et al. (2010). Native GABA_B_ receptors are heteromultimers with a family of auxiliary subunits. Nature 465, 231–235. 10.1038/nature08964.

Schwenk, J., Perez-Garci, E., Schneider, A., Kollewe, A., Gauthier-Kemper, A., Fritzius, T., Raveh, A., Dinamarca, M.C., Hanuschkin, A., Bildl, W., et al. (2016). Modular composition and dynamics of native GABAB receptors identified by high-resolution proteomics. Nat Neurosci 19, 233–242. 10.1038/nn.4198.

Sclip, A., and Sudhof, T.C. (2023). Combinatorial expression of neurexins and LAR-type phosphotyrosine phosphatase receptors instructs assembly of a cerebellar circuit. Nat Commun 14, 4976. 10.1038/s41467-023-40526-0.

Siddiqui, T.J., Pancaroglu, R., Kang, Y., Rooyakkers, A., and Craig, A.M. (2010). LRRTMs and neuroligins bind neurexins with a differential code to cooperate in glutamate synapse development. J Neurosci 30, 7495–7506. 10.1523/JNEUROSCI.0470-10.2010.

Soler-Llavina, G.J., Arstikaitis, P., Morishita, W., Ahmad, M., Sudhof, T.C., and Malenka, R.C. (2013). Leucine-rich repeat transmembrane proteins are essential for maintenance of long-term potentiation. Neuron 79, 439–446. 10.1016/j.neuron.2013.06.007.

Sterky, F.H., Trotter, J.H., Lee, S.J., Recktenwald, C.V., Du, X., Zhou, B., Zhou, P., Schwenk, J., Fakler, B., and Sudhof, T.C. (2017). Carbonic anhydrase-related protein CA10 is an evolutionarily conserved pan-neurexin ligand. Proc Natl Acad Sci U S A 114, E1253–E1262. 10.1073/pnas.1621321114.

Sudhof, T.C. (2017). Synaptic Neurexin Complexes: A Molecular Code for the Logic of Neural Circuits. Cell 171, 745–769. 10.1016/j.cell.2017.10.024.

Szoboszlay, M., Kirizs, T., and Nusser, Z. (2017). Objective quantification of nanoscale protein distributions. Sci Rep 7, 15240. 10.1038/s41598-017-15695-w.

Takahashi, H., and Craig, A.M. (2013). Protein tyrosine phosphatases PTPdelta, PTPsigma, and LAR: presynaptic hubs for synapse organization. Trends Neurosci 36, 522–534. 10.1016/j.tins.2013.06.002.

Takahashi, H., Katayama, K., Sohya, K., Miyamoto, H., Prasad, T., Matsumoto, Y., Ota, M., Yasuda, H., Tsumoto, T., Aruga, J., and Craig, A.M. (2012). Selective control of inhibitory synapse development by Slitrk3-PTPdelta trans-synaptic interaction. Nat Neurosci 15, 389–398, S381-382. 10.1038/nn.3040.

Tang, A.H., Chen, H., Li, T.P., Metzbower, S.R., MacGillavry, H.D., and Blanpied, T.A. (2016). A trans-synaptic nanocolumn aligns neurotransmitter release to receptors. Nature 536, 210–214. 10.1038/nature19058.

Trotter, J.H., Hao, J., Maxeiner, S., Tsetsenis, T., Liu, Z., Zhuang, X., and Sudhof, T.C. (2019). Synaptic neurexin-1 assembles into dynamically regulated active zone nanoclusters. J Cell Biol 218, 2677–2698. 10.1083/jcb.201812076.

Uemura, T., Lee, S.J., Yasumura, M., Takeuchi, T., Yoshida, T., Ra, M., Taguchi, R., Sakimura, K., and Mishina, M. (2010). Trans-synaptic interaction of GluRdelta2 and Neurexin through Cbln1 mediates synapse formation in the cerebellum. Cell 141, 1068–1079. 10.1016/j.cell.2010.04.035.

Um, J.W., Choi, T.Y., Kang, H., Cho, Y.S., Choii, G., Uvarov, P., Park, D., Jeong, D., Jeon, S., Lee, D., et al. (2016). LRRTM3 Regulates Excitatory Synapse Development through Alternative Splicing and Neurexin Binding. Cell Rep 14, 808–822. 10.1016/j.celrep.2015.12.081.

Ushkaryov, Y.A., Petrenko, A.G., Geppert, M., and Sudhof, T.C. (1992). Neurexins: synaptic cell surface proteins related to the alpha-latrotoxin receptor and laminin. Science 257, 50–56. 10.1126/science.1621094.

Van Rossum, G., and Drake Jr, F. (1995). Python reference manual (Centrum voor Wiskunde en Informatica Amsterdam).

Varoqueaux, F., Aramuni, G., Rawson, R.L., Mohrmann, R., Missler, M., Gottmann, K., Zhang, W., Sudhof, T.C., and Brose, N. (2006). Neuroligins determine synapse maturation and function. Neuron 51, 741–754. 10.1016/j.neuron.2006.09.003.

Wentzel, C., Sommer, J.E., Nair, R., Stiefvater, A., Sibarita, J.B., and Scheiffele, P. (2013). mSYD1A, a mammalian synapse-defective-1 protein, regulates synaptogenic signaling and vesicle docking. Neuron 78, 1012–1023. 10.1016/j.neuron.2013.05.010.

Wong, M.Y., Liu, C., Wang, S.S.H., Roquas, A.C.F., Fowler, S.C., and Kaeser, P.S. (2018). Liprin-alpha3 controls vesicle docking and exocytosis at the active zone of hippocampal synapses. Proc Natl Acad Sci U S A 115, 2234–2239. 10.1073/pnas.1719012115.

Woo, J., Kwon, S.K., Choi, S., Kim, S., Lee, J.R., Dunah, A.W., Sheng, M., and Kim, E. (2009). Trans-synaptic adhesion between NGL-3 and LAR regulates the formation of excitatory synapses. Nature Neuroscience 12, 428–437. 10.1038/nn.2279.

Yan, Q., Weyn-Vanhentenryck, S.M., Wu, J., Sloan, S.A., Zhang, Y., Chen, K., Wu, J.Q., Barres, B.A., and Zhang, C. (2015). Systematic discovery of regulated and conserved alternative exons in the mammalian brain reveals NMD modulating chromatin regulators. Proc Natl Acad Sci U S A 112, 3445–3450. 10.1073/pnas.1502849112.

Yuzaki, M. (2018). Two Classes of Secreted Synaptic Organizers in the Central Nervous System. Annu Rev Physiol 80, 243–262. 10.1146/annurev-physiol-021317-121322.

Zhang, P., Lu, H., Peixoto, R.T., Pines, M.K., Ge, Y., Oku, S., Siddiqui, T.J., Xie, Y., Wu, W., Archer-Hartmann, S., et al. (2018). Heparan Sulfate Organizes Neuronal Synapses through Neurexin Partnerships. Cell 174, 1450–1464 e1423. 10.1016/j.cell.2018.07.002.

